# Closing the biodiversity observation-to-action loop

**DOI:** 10.64898/2026.08.27.747669

**Authors:** Keiji Yamaguchi, Kei Uchida, Masayoshi K. Hiraiwa, Yuya Fukano

## Abstract

Citizen science observations are abundant, but conservation requires turning uneven records into reliable predictions and directing new surveys to where information is missing. We developed a biodiversity platform for Japan that is updated monthly and integrates 2.32 million records to predict 8,297 species across seven taxonomic groups. Shared representation models outperformed species-specific models in four groups and extended predictions to species with few records. Five independent datasets, including structured monitoring, environmental DNA and complete forest inventories, confirmed that the models ranked observed species and occupied sites above alternatives, with median AUCs of 0.724 to 0.894 across sites and 0.650 to 0.841 across species. For any user-selected area, the platform returns candidate species, distribution predictions, a biodiversity map corrected for uneven observation effort, a conservation priority map for native species and a map recommending where to survey next. This map highlights places where species with few records are predicted to occur despite limited sampling. Independent observations showed that areas ranked highly by this predicted potential contained many such species, indicating that model predictions can help direct surveys toward knowledge gaps. New observations are incorporated into monthly updates, creating a national feedback system connecting citizen science, local conservation decisions and future surveys.

## 1. Introduction

Reversing biodiversity loss requires not only international targets and protected areas, but also a system that continuously monitors biodiversity status in each region and rapidly feeds this information back into land management, conservation planning, and further surveys (CBD, 2022; Danielsen et al., 2024). This need is explicitly recognized in Target 21 of the Kunming–Montreal Global Biodiversity Framework, which calls for the best available biodiversity data, information and knowledge to be made accessible to decision makers, practitioners and the public to guide effective biodiversity action (CBD, 2022). Because the distribution of biodiversity and its changes are spatially and temporally heterogeneous, systematic surveys conducted by a limited number of experts alone cannot continuously assess biodiversity status across broad areas and multiple taxonomic groups. Citizen science has become an important means of filling these observation gaps by accumulating large numbers of biodiversity observations across spatial and temporal ranges that are difficult to cover through conventional surveys (Chandler et al., 2017; Pocock et al., 2018). As such observations accumulate, the emerging challenge is no longer simply to collect more biodiversity records, but to transform uneven data into reliable, locally meaningful and decision-relevant knowledge. Citizen observations must be converted into spatial predictions, their reliability and limitations must be evaluated, and the resulting information must be returned for use in local conservation decision-making. These outputs must also be linked to subsequent field verification and further observations, thereby creating a cyclical information infrastructure that connects observation, prediction, validation, and decision-making.

Species distribution models (SDMs) play a central role in this cycle by converting heterogeneous occurrence records into spatially explicit predictions of habitat suitability and species distributions (Elith and Leathwick, 2009; Guisan et al., 2013). Recent advances include deep-learning models that learn many species simultaneously and systems that continuously incorporate new observation data into predictions. For example, Brun et al. (2024) used approximately 6.7 million citizen-science records of 2,477 plant species in Switzerland and showed that a multi-species deep neural network improved predictions of species distributions and community composition. Ovaskainen et al. (2026) integrated smartphone-based bird-sound observations, long-term monitoring, and detection models to implement a biodiversity digital twin that is continuously updated with new citizen-science data. More broadly, BON in a Box provides open workflows for transforming biodiversity data into indicators and identifies data-gap assessment and spatial sampling prioritization as central components of biodiversity monitoring (Griffith et al., 2026). Together, these developments show that many of the technical components needed to connect biodiversity observations, prediction, monitoring and decision support are already available. The remaining challenge is to integrate and evaluate these components as an operational national-scale system that extends across multiple taxonomic groups and data-poor species, establishes reliability through independent validation, and links the resulting information to local use and further observation.

First, an information platform for local conservation must cover not only a small number of well-observed species but also multiple taxonomic groups and many data-poor species. Existing large-scale prediction systems have mainly focused on a single taxonomic group with relatively rich occurrence records, such as plants or birds (Fink et al., 2020; Brun et al., 2024; Ovaskainen et al., 2026). However, local biodiversity includes plants, vertebrates, invertebrates, and aquatic organisms, and a single taxonomic group cannot fully represent local conservation value or survey needs. Public occurrence data also form a taxonomic long tail, in which many records are concentrated in a small number of frequently observed species, whereas most species have only a few records (Troudet et al., 2017; Johnston et al., 2023; Bowler et al., 2025). Here, data-poor species are defined as species with few occurrence records and are not necessarily species with low abundance or threatened species. Multi-species models that jointly learn many species in a shared latent representation and share information on environmental responses among species may reduce this imbalance (Chen et al., 2017; Tikhonov et al., 2020; Pichler and Hartig, 2021; Brun et al., 2024). Indeed, information shared among species in community data can improve distribution predictions for species with few records (Zhang et al., 2020). However, it remains unclear whether shared learning can extend prediction across a broad range of taxonomic groups and expand the number of data-poor species that can be included in national biodiversity predictions. Graph neural networks that incorporate the spatial structure of landscapes and sampling locations have also recently been introduced into SDMs (Wu et al., 2025), but it remains unclear whether the gains from multi-species learning arise from spatial graph structure or from simpler shared representations among species.

Second, using predictions based on citizen science for public decision-making for biodiversity conservation requires validation beyond the internal evaluation used during model development. With presence-only data, failure to record a species may indicate true absence, non-detection, or lack of survey, and evaluation metrics are strongly affected by the selection of background sites and bias in observation effort (Lobo et al., 2008; Lahoz-Monfort et al., 2014; Guillera-Arroita et al., 2015). Randomly dividing spatially close records between training and evaluation data can also overestimate predictive performance in new regions because of spatial autocorrelation (Roberts et al., 2017). Moreover, relative habitat-suitability scores can be misinterpreted as absolute probabilities of occurrence or confirmed species lists, and maps created by stacking predictions across species may retain geographic bias in observation effort. Public information platforms for conservation therefore require rigorous spatial partitioning and independent validation using data collected through observation processes different from those used for model training. Although stacked biodiversity maps are useful for comparing regions and identifying priority areas, errors in individual species models and bias in observation effort can propagate into estimates of species richness and community composition and distort apparent diversity patterns (Calabrese et al., 2014; Scherrer et al., 2020; Zwiener and Alves, 2023; Baker et al., 2024). These effects must be evaluated and clearly reported, and the resulting maps should be presented as indicators for further surveys and field verification rather than as definitive estimates of species richness or conservation priority.

Third, predictions based on citizen science must be returned to local users and linked to further observations. Adaptive sampling can improve models by directing surveys to sites with high information value and adding new records to model training. Simulations suggest that directing some observers to sites with high model uncertainty can improve SDM performance (Mondain-Monval et al., 2024). In practice, however, citizen scientists often select not only accessible sites but also known natural areas and species-rich locations, so existing observation behaviour may already be an effective search strategy (Tulloch and Szabo, 2012; Arazy and Malkinson, 2021; Dimson and Gillespie, 2023). Moreover, sites with high uncertainty, few records, and high predicted habitat suitability for data-poor species may differ. External data are therefore needed to test whether model-based survey prioritization improves average predictive performance or instead redirects effort towards data-poor species and regions, thereby expanding taxonomic coverage.

In this study, we developed an open biodiversity intelligence platform that converts nationwide citizen-science observations into externally validated, multi-taxon predictions for local conservation and links these predictions to further observations. Across Japan, we integrated 2.32 million occurrence records from GBIF, iNaturalist, eBird, the Ministry of the Environment’s Ikimono Log, and researcher surveys with national environmental data on topography, climate, land use, distances to landscape features, soils, and satellite observations. We predicted the distributions of 8,297 species across seven major taxonomic groups. For any user-defined area, the platform provides existing records, predicted candidate and red-listed species, observation coverage, stacked predictions, observation effort, and survey-priority areas.

Within this operational framework, we addressed three scientific questions about converting heterogeneous citizen-science observations into reliable and actionable biodiversity information at a national scale. First, we tested across taxonomic groups and training scales whether shared-representation models outperformed species-specific models and whether including data-poor species improved predictions for well-sampled species or expanded predictive coverage. Second, beyond spatial block partitioning, we evaluated predictions using five external datasets collected through different observation processes: species lists obtained from certification materials for 393 Nationally Certified Sustainably Managed Natural Sites (hereafter, certified natural sites) recognized under Japan’s 30by30 initiative, environmental DNA records from 148 ANEMONE MiFish sites, structured survey records from 109 Monitoring Sites 1000 sites, complete stem-census records from 60 forest plots, and plot-level tree records from the Japanese National Forest Inventory. We also used geographically separated holdout data and external presence–absence data to assess and calibrate predicted occurrence frequencies, and tested whether stacked biodiversity maps retained geographic bias in observation effort. Third, we compared survey-priority strategies based on predictive uncertainty, observation gaps, existing observation effort, and predicted habitat suitability for data-poor species, and evaluated their effects on average predictive performance and the discovery of data-poor species. Together, these analyses tested a framework for converting citizen-science observations into transparent and verifiable biodiversity information that supports local conservation planning and guides further observations.

## 2. Results

### 2.1. Integration of occurrence data and development of the national prediction platform

We developed the National Adaptive Nature Assessment and Knowledge Updating System for Action (NANAKUSA), a biodiversity information platform that converts citizen-science and publicly available occurrence data into nationwide multi-species predictions and returns the results to local users (Fig. 1; URL: https://mapryam.net/). We collected and integrated occurrence records from GBIF, iNaturalist, eBird, the Ministry of the Environment’s Ikimono Log, and field surveys conducted by researchers. After deduplication, the master dataset contained 2,322,578 records for 37,995 species from 1990 onward. Of these, 1,461,834 records for 26,869 species fell within the 2020– 2025 observation window. We initially selected 5,608 species with at least 20 records for model training and applied long-tail pooling to fish, arthropods, and plants and fungi by lowering the minimum threshold to five records. At deployment, the shared-representation models had output heads for 7,987 species: 451 birds, 480 fish, 3,578 arthropods, 151 other invertebrates, and 3,327 plants and fungi. We linked these records to 41 nominal environmental covariates describing topography, climate, land use, distances to landscape features, soils, and satellite observations, of which approximately 29 were effectively available for modelling (Supplementary Methods 1; Supplementary Tables 1 and 2). The number of occurrence records was highly uneven among species, forming a long-tailed distribution in which a small number of species had many records and most species had few records (Supplementary Fig. 1).

**Figure 1.**
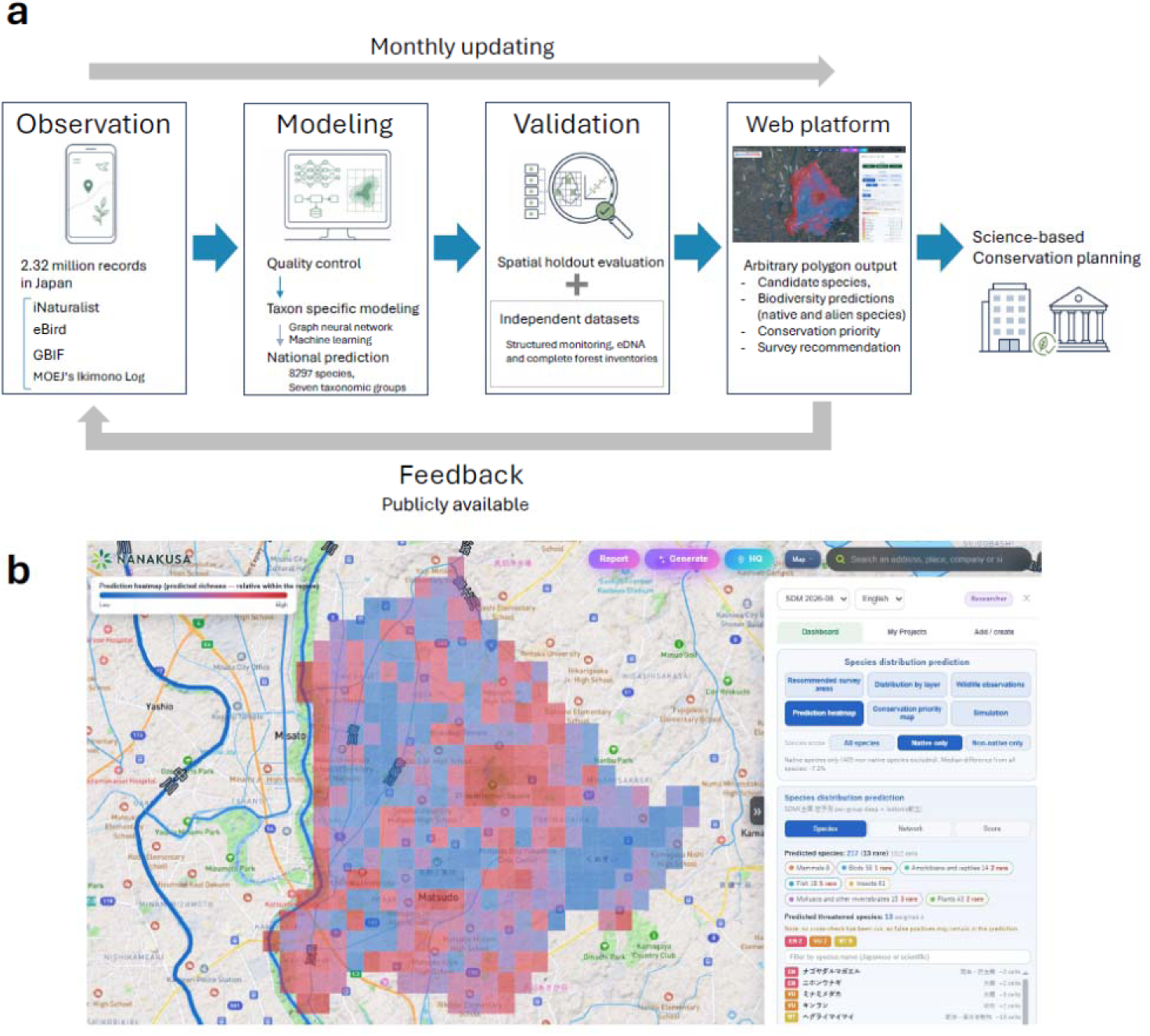
Nationwide biodiversity information cycle and an example output from the web platform. **a,** Occurrence records from multiple sources undergo quality control and taxon-specific modelling to generate nationwide predictions for 8,297 species across seven taxonomic groups. Predictions are evaluated using spatial holdouts and independent datasets before being delivered through the web platform as candidate-species lists, biodiversity predictions, conservation priorities and survey recommendations. New publicly available observations are incorporated into monthly updates, creating a feedback cycle from observation to conservation planning and back to observation. **b,** Example of the native-species prediction heatmap for Matsudo City, Chiba Prefecture, Japan. Grid-cell colours indicate relative predicted native-species richness within the selected area, from lower values in blue to higher values in red, and do not represent absolute species richness.

Based on the performance evaluation described below, we used shared-representation deep models for birds, fish, arthropods, other invertebrates, and plants and fungi, and species-specific ensemble models for mammals and reptiles and amphibians. We also applied long-tail pooling to fish, arthropods, and plants and fungi by adding data-poor species to shared training. Species selection, spatial partitioning, background-site sampling, model architecture, hyperparameters, and taxon-specific deployment rules are described in Supplementary Methods 1 and Supplementary Tables 1 and 3. This procedure produced nationwide predictions for 8,297 species. Among the 8,290 species entering the stacked layers (the 8,297 deployed species less seven human and domesticated taxa; Methods), 542 were classified as alien using a curated GRIIS-based classification. These species accounted for 7.10% of the training records and 7.97% of the nationwide stacked prediction when all deployed species were included, had approximately 3.5 times more records per species than native species, and were concentrated in urban areas. Prediction surfaces for each species were provided on a 20-m output grid, allowing users to obtain species-level predictions and regional summaries for any selected location or area.

The platform was implemented as an updating system rather than a static prediction map. Each month, it retrieves new occurrence records, performs taxonomic and coordinate quality control, updates the training data, retrains taxon-specific models, recalculates national prediction surfaces and regional indicators, and redeploys the outputs to the web application. Candidate species, stacked predictions, observation effort, and survey-priority areas are therefore updated as new observations accumulate. Each public release records the cut-off date of the occurrence data, the model version, and the update date. The taxonomic group, number of occurrence records, and selected model for all deployed species are provided in Supplementary Data 1.

### 2.2. Performance of shared-representation models

Under the same spatial holdout design, shared-representation deep models showed higher discrimination than species-specific random forests in four of the seven taxonomic groups (Fig. 2a). The median difference in species-level ROC-AUC was +0.025 for birds, +0.040 for fish, +0.078 for arthropods, and +0.148 for other invertebrates. The differences remained significant after correction for multiple comparisons in birds and arthropods. Improvements were larger for species with lower performance under species-specific models. In contrast, shared-representation models showed no clear advantage for mammals, reptiles and amphibians, or plants and fungi. Robustness analyses using different random seeds and alternative species-specific model settings produced broadly similar taxonomic patterns (Supplementary Methods 2; Supplementary Fig. 2; Supplementary Table 4). Non-spatial random partitioning substantially inflated estimated performance relative to spatial block partitioning. Among 554 species evaluated under both designs, 91% had equal or higher species-level ROC-AUC under random partitioning, and the median increased from 0.634 to 0.858 (Fig. 3b).

**Figure 2.**
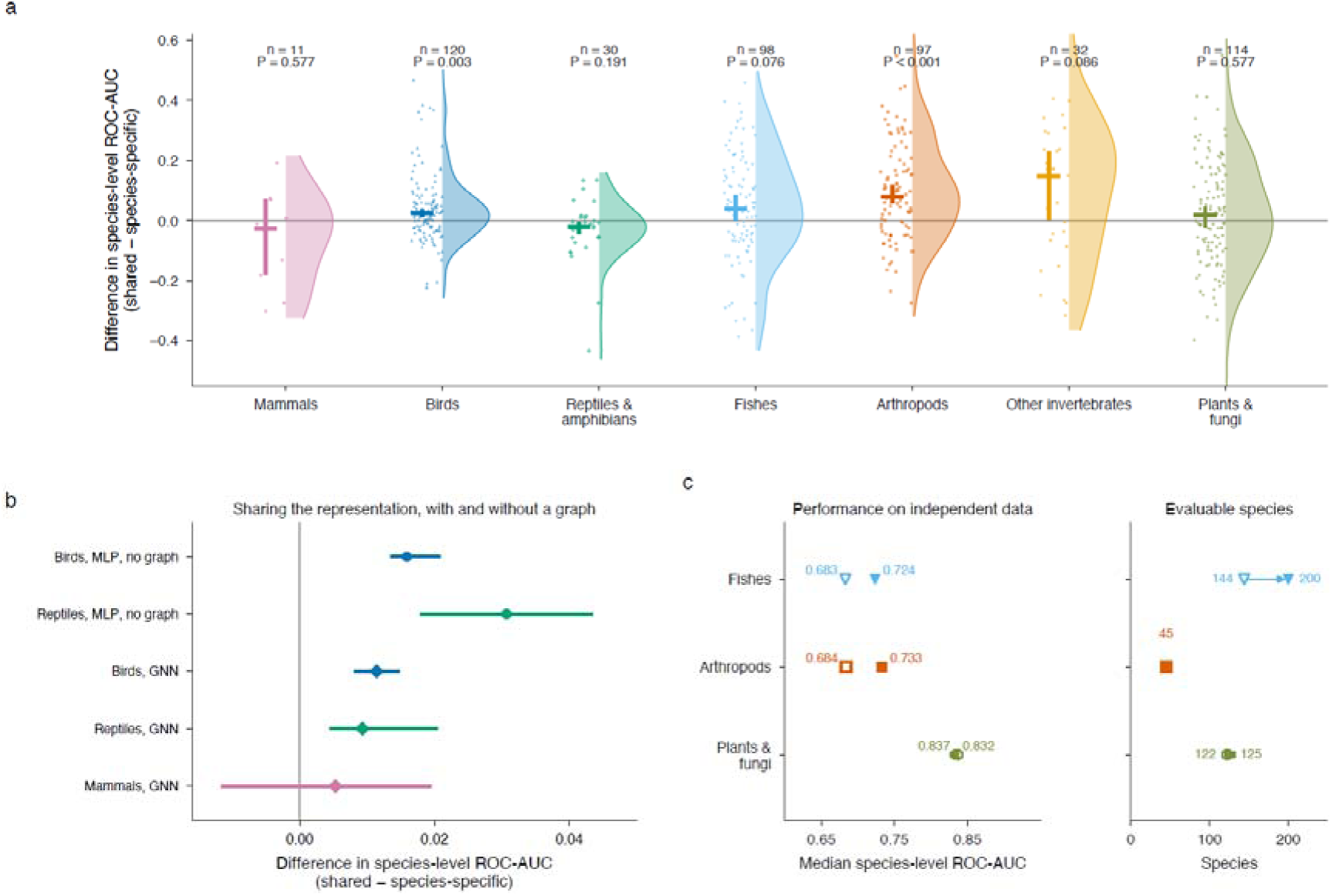
Shared representations improve multispecies prediction and extend taxonomic coverage. a, Differences in species-level ROC-AUC between shared-representation and species-specific models under the same spatial holdout design. Points represent species, half-violins show the distribution within each taxonomic group and positive values indicate higher discrimination by the shared model. Sample sizes and P values are shown above each group; P values are Holm-adjusted across the seven taxonomic groups. b, Ablation comparisons of shared and species-specific models using graph-free multilayer perceptrons (MLPs) and graph neural networks (GNNs). Points and horizontal intervals summarize the differences in species-level ROC-AUC. c, Independent-data performance and numbers of evaluable species before long-tail pooling (open symbols) and after pooling (filled symbols). Because species were not filtered by native or domestication status (Methods 4.2), the mammal panel includes human, domesticated, alien and marine species (7 of the 11 evaluable species). Restricting the panel to the four native wild mammals yields a median ΔAUC of −0.030, consistent with the full-panel median of −0.027; the choice of per-species ensembles for mammals is unaffected.

**Figure 3.**
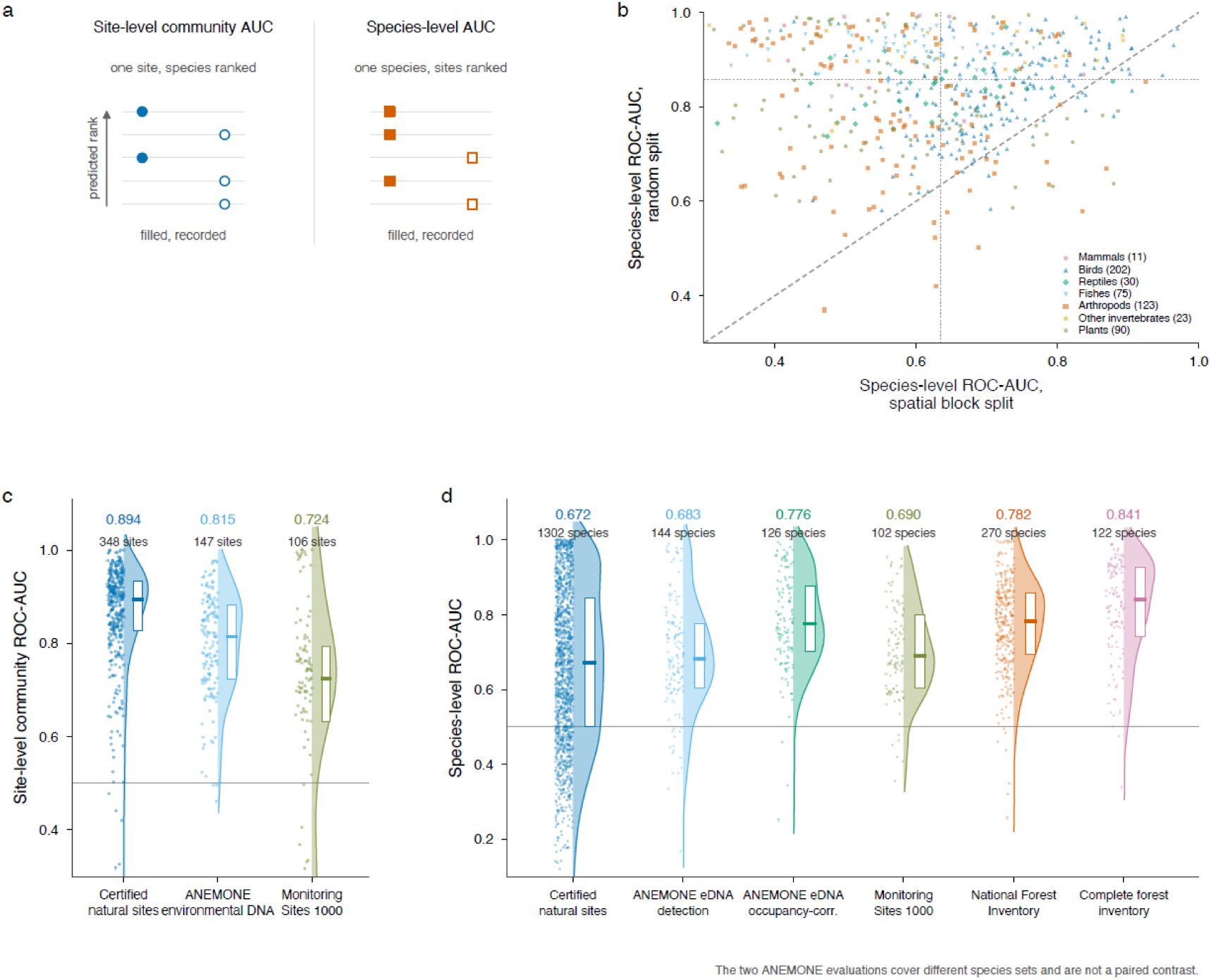
Internal and external evaluation of predictive discrimination. **a,** Schematic definitions of site-level community and species-level ROC-AUC. Site-level community AUC ranks species within a site, whereas species-level AUC ranks sites for a species. Filled symbols denote species recorded at the focal site in the site-level schematic and sites where the focal species was recorded in the species-level schematic; open symbols denote the corresponding non-recorded species or sites. **b,** Species-level ROC-AUC under spatial-block and non-spatial random splits for 554 species evaluable under both designs. The diagonal indicates equal performance. **c,** Site-level community ROC-AUC for certified natural sites (n = 348 evaluable sites), ANEMONE environmental DNA (n = 147) and Monitoring Sites 1000 (n = 106). Points represent sites, half-violins show the distribution, boxes show the interquartile range and central lines show medians; the median and the number of evaluable sites are also printed above each distribution. **d,** Species-level ROC-AUC for six external evaluations, from left to right: certified natural sites (unadjusted evaluation; n = 1,302 species), ANEMONE eDNA scored with detection–non-detection labels (detection; n = 144), the occupancy-corrected ANEMONE evaluation (occupancy-corr.; n = 126), Monitoring Sites 1000 (n = 102), the National Forest Inventory (n = 270) and complete forest inventories (n = 122). Points represent species, half-violins show the distribution, boxes show the interquartile range and central lines show medians; the median and the number of evaluable species are also printed above each distribution. The two ANEMONE evaluations cover different species sets and are not a paired contrast; the 98 species represented in both are compared in Supplementary Fig. 4e.

We next compared the full deployment pipelines using an independent complete forest inventory of 125 tree species. The median species-level ROC-AUC was 0.832 for the shared-representation model, 0.806 for MaxEnt, and 0.761 for the species-specific random forest, with the shared-representation model significantly outperforming both alternatives (Supplementary Fig. 2d). However, the sets of environmental covariates differed among these pipelines. We therefore conducted a controlled comparison for 28 species using the same occurrence records, background sites, environmental covariates, and spatial partitioning. Under these conditions, all models performed near chance level, and MaxEnt and random forests performed similarly to or slightly better than the deep-learning models (Supplementary Fig. 2; Supplementary Table 4).

Independent validation showed that long-tail pooling yielded informative predictions for some additional low-record species, rather than merely increasing the number of prediction targets. Here, an “externally evaluable species” was a species for which the independent dataset met the dataset-specific requirements for calculating ROC-AUC (Supplementary Methods 2). For fish, the median species-level ROC-AUC increased from 0.68 to 0.72, and the number of evaluable species increased from 144 to 200. The 56 newly evaluable fish species had a median ROC-AUC of 0.715. Median ROC-AUC increased from 0.68 to 0.73 for arthropods. For plants and fungi the number of externally evaluable species increased from 122 to 125; on the 122 species and 59 inventory plots evaluable in both arms, pooling left discrimination essentially unchanged (median calibrated AUC 0.837 before and 0.834 after; paired median difference +0.001, 95% CI -0.009 to +0.009, P = 0.52; Supplementary Table 4). Performance among the newly added data-poor species varied strongly among taxonomic groups, and some species remained near chance level (Supplementary Methods 2; Supplementary Fig. 3; Supplementary Table 4).

To assess the contribution of the spatial graph, we compared a graph-free shared model, which learned a common environmental representation across species without exchanging information among locations, with a graph-based shared model that additionally propagated information among locations connected in the spatial graph. Ablation analyses showed that the gain from sharing the representation was at least as large without a spatial graph as with one, and that neither reweighting or densifying the neighbourhood graph nor adding river- or catchment-based connections improved median ROC-AUC by more than 0.01. In reptiles and amphibians, the only group in which the graph-based and graph-free shared models were evaluated on identical data partitions, the shared graph neural network scored slightly higher than the shared multilayer perceptron (paired median difference +0.016, 95% CI +0.001 to +0.035, P = 0.0015, 18 of 25 species); because the two runs were not matched on hyperparameter tuning or training epochs, this difference cannot be attributed to the spatial graph itself, and the shared model did not exceed the species-specific random-forest baseline (+0.009, P = 0.46). These results indicate that the main benefit of the shared-representation models arose from learning environmental representations across species; the additional contribution of propagating spatial information among sites through a graph was small where it could be isolated (Fig. 2b; Supplementary Methods 2; Supplementary Fig. 3; Supplementary Table 4).

### 2.3. External validation of model predictions

We evaluated the nationally deployed models using five datasets with observation designs and detection processes different from those of the training data: certified natural sites, Monitoring Sites 1000, the National Forest Inventory, ANEMONE environmental DNA surveys, and complete forest inventories (Supplementary Methods 3). We calculated site-level community AUC, which measures whether recorded species were ranked above unrecorded species within each site, and species-level AUC, which measures whether recorded sites were ranked above unrecorded sites for each species (Supplementary Methods 3; Supplementary Table 5).

For the certified natural sites, we matched 1,302 species across 393 sites. Among 348 evaluable sites, the median site-level community AUC was 0.894, and recorded species were ranked within the top 16.3% of candidate species from the same taxonomic group (Fig. 3c). After excluding site–species combinations with training records of the same species within 2 km of the site boundary, the median remained 0.871 across 335 sites. For birds, retraining after removing nearby records changed AUC by only 0.002–0.006. The median species-level AUC was 0.672 and decreased to 0.650 after controlling for nearby records. Among 127 data-poor species added through long-tail pooling, the median was 0.500, and 49.6% had AUC values below 0.5 (Supplementary Fig. 4; Supplementary Table 5).

The models also performed above chance in two standardized, broad-scale survey datasets. For Monitoring Sites 1000, which included 109 sites and 102 species, the median site-level community AUC was 0.724 across the 106 evaluable sites and the median species-level AUC was 0.690. In the National Forest Inventory, which included 15,835 plots and 270 tree species, the median species-level AUC was 0.782, and 73.7% of species had AUC values above 0.7 (Fig. 3c, d; Supplementary Fig. 4; Supplementary Table 5).

Two additional datasets allowed non-records to be treated more explicitly. The ANEMONE dataset included repeated environmental DNA surveys, allowing imperfect detection to be modelled using occupancy models. The dataset comprised 148 sites and 144 fish species. The median site-level community AUC was 0.815 across the 147 evaluable sites, and the median species-level AUC, computed from detection–non-detection labels, was 0.683 across the 144 species (ANEMONE eDNA, detection, in Fig. 3d). In a partly overlapping set of 126 species used for the occupancy analysis (occupancy fitted across 137 sites; AUC evaluated at the 123 sites with available predictions), the median species-level AUC, calculated from the same detection–non-detection labels, was 0.776 (ANEMONE eDNA, occupancy-corrected, in Fig. 3d). The two ANEMONE evaluations cover different species sets and are not a paired contrast (Supplementary Fig. 4e; Supplementary Table 5). The complete forest inventories recorded all trees within 60 plots, allowing unrecorded tree species to be treated as absences. Across 122 tree species, the median species-level AUC was 0.841, the highest among the external datasets.

These results show that species-level evaluation depended strongly on the observation process and the meaning of non-records, and performance was highest when non-records could be treated as confirmed absences.

### 2.4. Calibration of prediction scores

Separate from the nationwide deployment, we tested the geographic transferability of isotonic calibration for four taxonomic groups in central Japan. For an ensemble averaging random-forest and logistic-regression predictions, we fitted calibration functions using the western third of the study region, excluded the central third as a buffer, and evaluated calibration in the eastern third (Supplementary Methods 4).

Calibration largely preserved the ranking of predictions while reducing expected calibration error from 0.16–0.28 to below 0.003 and Brier score from 0.064–0.117 to 0.014–0.029 (Fig. 4a; Supplementary Fig. 5; Supplementary Table 6). For nationwide deployment, calibration models corresponding to each taxonomic group and model pathway were applied.

**Figure. 4.**
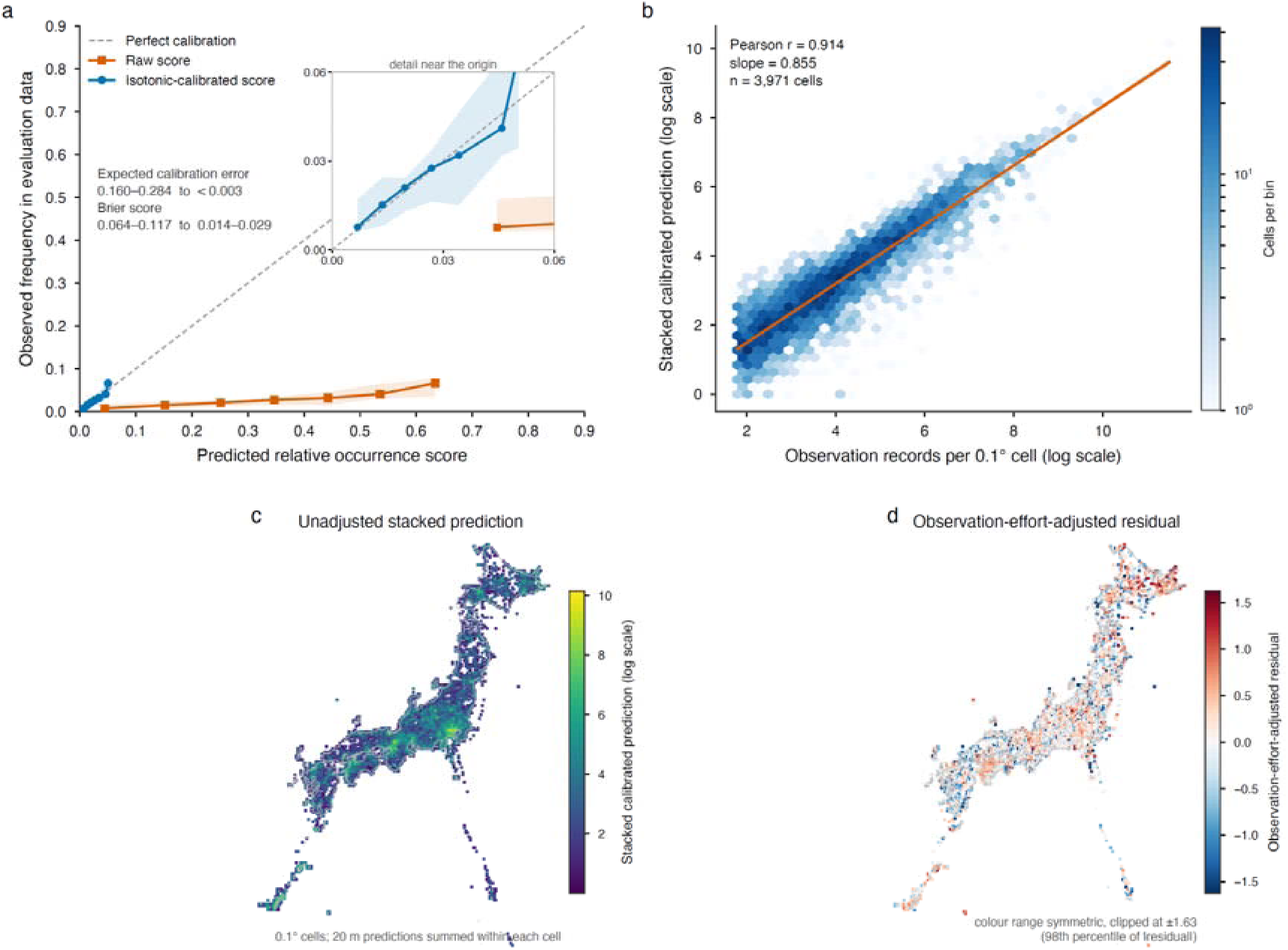
Calibration and observation effort shape stacked biodiversity predictions. **a,** Reliability of raw and isotonic-calibrated relative occurrence scores. Lines show count-weighted aggregates across four taxonomic groups, shading shows the range among groups and the diagonal represents perfect calibration. The two highest raw-score bins, containing 0.11% of evaluation points, were omitted. Calibration reduced expected calibration error from 0.160– 0.284 to below 0.003 and Brier score from 0.064–0.117 to 0.014–0.029. **b,** Relationship between observation records and the stacked calibrated prediction across 3,971 0.1° cells. Hexagon colour represents the number of cells and the orange line shows the fitted relationship (r = 0.914, slope = 0.855). **c,** National unadjusted stacked prediction. **d,** Residual stacked prediction after accounting for observation effort. Panels b–d use calibrated predictions for 7,739 native species after additionally excluding nine marine mammals. The mapped values represent relative predicted biodiversity potential, not absolute species richness.

The occurrence frequencies used in this analysis were relative frequencies within datasets consisting of occurrence records and evaluation background sites and therefore did not represent absolute occurrence probabilities in nature. We therefore fitted separate calibration functions using presence–absence data from ANEMONE environmental DNA surveys for fish and complete forest inventories for plants. These external calibrations reduced expected calibration error to 0.026–0.035 (Supplementary Fig. 5; Supplementary Table 6). Thus, raw model outputs should not be interpreted as absolute occurrence probabilities, but they can be converted into scores corresponding to observed occurrence frequencies when calibration data with an appropriate observation design are available.

### 2.5. Relationship between stacked predictions and observation effort

We tested whether spatial bias in citizen-science observation effort remained after calibrated predictions were stacked across native species. Japan was divided into 0.1° grid cells, and calibrated scores were summed within each cell across the 7,739 native species that remained after seven human and domesticated taxa, 542 species classified as alien and nine marine mammals were removed, to obtain a stacked prediction value (Supplementary Methods 5). Stacked prediction values were strongly correlated with the log number of occurrence records. Across 3,971 cells containing at least five records, the correlation coefficient was 0.914 and the regression slope was 0.855 (Fig. 4b). High uncorrected values were concentrated in regions with many existing records, including western Tokyo and the Osaka–Kyoto metropolitan area (Fig. 4c). Thus, calibration of species-level scores did not remove geographic variation in observation effort from stacked biodiversity predictions.

To separate this effect, we regressed stacked prediction values against the number of occurrence records and calculated residuals from the values expected under similar observation effort. Areas with high residual values shifted away from major metropolitan regions towards Yakushima and eastern Hokkaido (Fig. 4d). These results show that stacked predictions based on citizen-science data should not be interpreted directly as differences in biodiversity without explicitly evaluating or correcting for observation effort.

### 2.6. Evaluation of survey-priority strategies

We first tested whether additional records from prioritized sites improved model performance more efficiently than randomly selected records. We removed part of the existing training data and then progressively returned records selected using different site-prioritization strategies (Supplementary Methods 6). Neither predictive uncertainty nor a composite score combining uncertainty, low observation effort, and data-poor-species potential restored species-level ROC-AUC on the fixed holdout data faster than random selection (Fig. 5a; Supplementary Fig. 6). Thus, prioritizing sites using uncertainty or the composite score did not efficiently improve average discrimination across species.

**Fig. 5.**
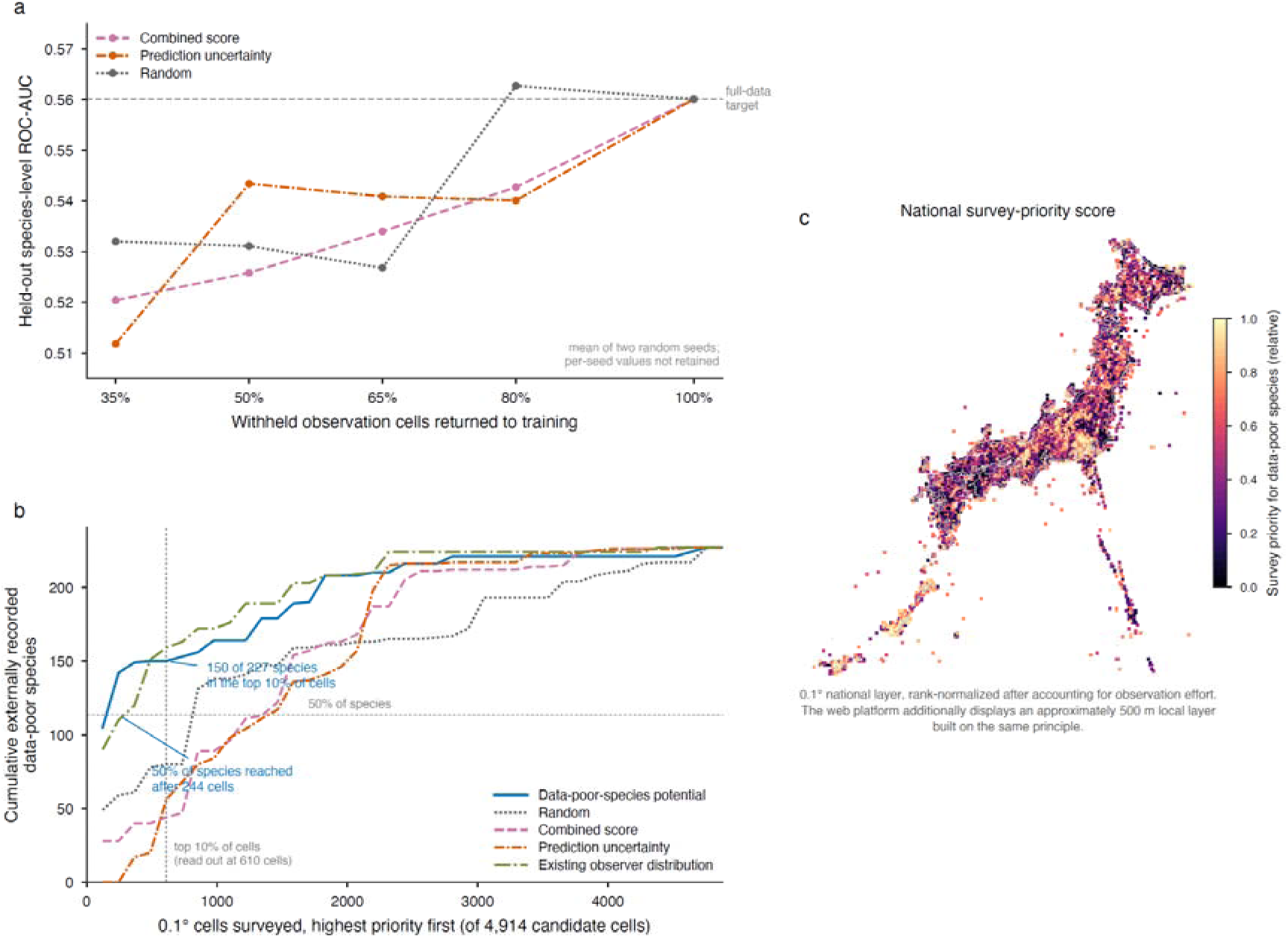
Survey-priority strategies differ between improving models and discovering data-poor species. **a**, Recovery of held-out species-level ROC-AUC as withheld observation cells were returned to training according to a combined score, prediction uncertainty or random selection. Lines show means from two random seeds; individual seed values were not retained. Neither model-based strategy recovered average discrimination faster than random selection. **b**, Cumulative number of externally recorded data-poor species detected when 4,914 candidate 0.1 ° cells were surveyed in descending priority order. The analysis compared data-poor-species potential, random selection, the combined score, prediction uncertainty and the existing observer distribution. Data-poor-species potential detected 150 of 227 species at the plotted 610-cell budget, the first evaluated budget point at or above the exact top decile of 491 cells; cumulative curves were evaluated every 122 cells. It reached 50% of the target species after 244 cells. **c,** National survey-priority score, rank-normalized after accounting for observation effort.

We next tested whether survey prioritization could improve the discovery of data-poor species using independent records from ANEMONE and Monitoring Sites 1000 that were not used for model training. Among 227 species with fewer than 20 training records, ranking candidate cells by uncorrected data-poor-species potential captured 150 species within the 610 highest-ranked cells and required 244 cells to capture 50% of the target species (Fig. 5b). The highest-ranked set contains 610 rather than the exact top decile of 491 of the 4,914 candidate cells because the cumulative curves were evaluated at 40 equal budget points (every 122 cells) and 610 was the first evaluated point at or above 491; the same set was used for every strategy. In comparison, uncertainty-based prioritization captured 56 species within the same 610 cells, and the composite score captured 44; both required 1,464 cells to capture 50% of the target species. Random selection captured 80 species within those cells and required 854 cells. Prioritization based on the existing distribution of observations captured 159 species within those cells and required 366 cells to capture 50%. Thus, data-poor-species potential captured half of the target species using the fewest cells, although it did not outperform the existing observation pattern within the 610-cell readout.

Because uncorrected data-poor-species potential was itself influenced by existing observation effort, the web application used residuals from a regression of this potential against the number of occurrence records as the survey-priority score. High values occurred mainly in natural areas of eastern Hokkaido, the Nansei Islands, and Kyushu (Fig. 5c; Supplementary Fig. 6). This score identifies areas where predicted signals for data-poor species exceed those expected from existing observation effort, rather than areas expected to maximize average model performance.

### 2.7. Implementation in a monthly updated web platform

We implemented the validated and calibrated nationwide predictions in a web platform that is updated monthly as new occurrence records accumulate (Fig. 1). The update pipeline retrieves new records from public data sources, performs taxonomic and coordinate quality control, rebuilds the training data, retrains taxon-specific models, recalibrates prediction scores, updates nationwide prediction surfaces and regional indicators, and redeploys the outputs to the web application. The platform therefore updates candidate-species lists, observation coverage, biodiversity predictions, and survey-priority areas rather than displaying a static snapshot. Users can select a prefecture, municipality, or registered site, or draw any polygon on the map. For the selected area, the platform displays existing records, predicted candidate species, species-level prediction maps, biodiversity predictions, conservation-priority maps, and survey-priority maps (Supplementary Methods 7; Supplementary Table 7). Existing records and model predictions are displayed separately, and each release reports the data cut-off date and model update date.

Species-level predictions remained available for all deployed species, including alien species. The biodiversity prediction represents stacked calibrated scores for the 8,290 species that remain after seven human and domesticated taxa are removed, after correction for observation effort; alien species are retained, and native-only and alien-only layers are served alongside the all-species layer. The conservation-priority map is restricted to native species and weights their calibrated scores by the inverse of each species’ nationwide mean calibrated score; including alien species would leave its ranking almost unchanged (Supplementary Methods 7.4). The survey-priority map uses residuals from the relationship between data-poor-species potential and observation effort. These regional indicators are displayed as relative values within the selected area and are intended for comparing candidate locations within that area, not for absolute comparisons among different areas. The conservation-priority map does not incorporate extinction risk, future land-use change, conservation cost, or management feasibility.

External validation assessed the uncorrected national data-poor-species potential at the 0.1° scale; the deployed residual score and the approximately 500-m local survey-priority surface were not directly validated.

## 3. Discussion

This study presents a framework for transforming nationwide citizen-science observations into a cyclical information infrastructure that supports both local conservation planning and future biodiversity surveys. The main contribution of the platform is not simply the visualization of multispecies predictions on a 20 m output grid. Rather, it integrates predictions across multiple taxonomic groups, external validation using datasets with different observation processes, calibration of model outputs, diagnosis of observation effort, delivery of information for user-defined regions, and identification of priority areas for additional surveys within a single public system.

This integration is implemented as a continuing operational process rather than a one-time analysis. New observations are incorporated each month, and the models, nationwide predictions and regional survey recommendations are then updated. The resulting platform is therefore not a static biodiversity map, but a publicly accessible information infrastructure that repeatedly updates both the current state of knowledge and survey priorities as new observations accumulate. Through this integration, our study makes several important contributions to the design and operation of conservation platforms based on nationwide biodiversity predictions.

First, our results show that a key value of shared-representation models for addressing the taxonomic long tail lies in expanding predictive coverage, in addition to improving prediction for some taxonomic groups. In analyses of the large nationwide dataset, shared-representation models outperformed species-specific models in several taxonomic groups (Fig2a), whereas their advantage was not clear in a smaller, fully controlled comparison (Supplementary Fig. 2c). The benefit therefore appears not to reflect a universal algorithmic advantage, but to emerge in large datasets that contain many species with highly uneven numbers of observations. Ablation analyses further indicated that this benefit arose mainly from environmental representations shared across species (Fig. 2b; Supplementary Fig. 3d,e); the additional contribution of the spatial graph was small in the one comparison where it could be isolated and could not be separated from differences in model configuration. By sharing information learned from well-observed species within taxonomic groups, shared-representation models may support predictions for species that are difficult to model separately. This property is particularly valuable for nationwide biodiversity platforms that must cover many species across multiple taxonomic groups (Tikhonov et al., 2020; Pichler and Hartig, 2021; Brun et al., 2024). Moreover, shared learning expanded predictive coverage towards the taxonomic long tail, rather than simply improving average predictive performance (Fig. 2c; Supplementary Fig. 3b,c). Adding data-poor species to the training set did not consistently improve performance for well-observed species, but it enabled some species that could not previously be modelled to be included as prediction targets. The main value of long-tail pooling therefore lies in expanding taxonomic coverage, rather than further increasing accuracy for species that were already predictable.

Second, this study goes beyond converting citizen-science observations into distribution predictions by establishing a nationwide operational cycle in which predictions guide future surveys and new records are returned to the models. Simulation studies have suggested that SDMs can be improved efficiently by directing observers to locations with high uncertainty or information value (Reich et al., 2018; Lange et al., 2023; Mondain-Monval et al., 2024). In our analysis, however, strategies based on prediction uncertainty or a composite score did not improve the ability of the models to distinguish occupied from unoccupied sites faster than random site selection (Fig. 5). One possible explanation is that citizen scientists with field experience and local knowledge already select locations where species are likely to be found, making current observation patterns a relatively effective search strategy. By contrast, prioritizing locations predicted to contain many data-poor species improved survey efficiency, capturing half of the target species using fewer sites than any of the other strategies (Fig. 5). The platform therefore presents survey-priority areas using residuals from the relationship between data-poor-species potential and existing observation effort. If new records accumulate in these areas and are incorporated into monthly model updates, the range of data-poor species that can be predicted and independently evaluated should continue to expand.

The same principle applies to biodiversity maps created by stacking predictions across many species. Although stacked maps can help identify relative biodiversity patterns and candidate survey areas within a region, they can also propagate observation biases shared across species-level models to the community level (Calabrese et al., 2014; Scherrer et al., 2020; Zwiener and Alves, 2023). In this study, stacked predictions remained strongly associated with observation effort after the calibrated scores of native species were combined (Fig. 4b). We therefore regressed the stacked native-species predictions against the number of observation records and presented the residuals. This index does not represent absolute species richness or a common scale that can be compared across regions. Instead, it shows the relative strength of the predicted biodiversity signal within the selected region that cannot be explained by observation effort alone. It should therefore be used not as evidence of the presence or absence of individual species or as a complete regional inventory, but as a hypothesis for checking existing inventories and narrowing the set of candidate locations for field surveys.

Third, this study shows that the reliability of citizen-science-based predictions for public use cannot be judged from a single evaluation design or accuracy metric. Random splitting produced substantially higher species-level AUCs than spatial-block splitting. Among the 554 species evaluated under both designs, random splitting produced equal or higher values for 91% of species, and the median AUC increased from 0.634 to 0.858 (Fig. 3). This result confirms that using spatially close training and evaluation sites can lead to overly optimistic estimates of predictive performance in unsampled areas (Hijmans, 2012; Roberts et al., 2017; Huang et al., 2025). External evaluation results also varied with the meaning of non-records and the evaluation method. In certified natural sites and structured monitoring datasets, non-records could not be clearly distinguished among true absences, non-detections and unsurveyed species. In ANEMONE, the occupancy-corrected evaluation that accounted for imperfect detection produced a higher species-level AUC. Complete forest inventories, in which non-records could be treated as absences, produced the highest species-level AUC among the external datasets (Fig. 3). At certified natural sites, the models performed well in ranking recorded species near the top of the candidate list for each site, but were less effective at distinguishing recorded from non-recorded sites for individual species, with substantial variation among species. Evaluation results are therefore not properties of the model alone, but depend on the spatial partitioning design, detection process, definition of non-records, evaluation unit and choice of metric (Lobo et al., 2008; Lahoz-Monfort et al., 2014; Guillera-Arroita et al., 2015; Roberts et al., 2017). Public biodiversity information infrastructures should therefore report multiple evaluations that match their intended uses, rather than presenting only the highest accuracy value, and should clearly explain what each evaluation tests. As one example, the high site-level community AUC obtained for certified natural sites supports the use of predicted candidate-species lists as a starting point for checking existing inventories and planning field surveys.

Probability calibration is especially important when species-level predictions are stacked. Major S-SDM frameworks have focused on whether continuous outputs should be summed, converted into binary predictions or constrained by community-level richness, rather than on explicitly calibrating component scores to observed frequencies (Calabrese et al., 2014; Scherrer et al., 2020; Zwiener and Alves, 2023). Yet summing continuous outputs has an expected-richness interpretation only if those outputs are calibrated occurrence probabilities, and raw machine-learning scores should not be averaged probabilistically without checking this correspondence (Dormann, 2020). In our analysis, isotonic calibration substantially improved agreement between model scores and observed frequencies, but calibration based on presence–background data remained conditional on that observation design and did not identify absolute occupancy probabilities (Phillips and Elith, 2013). External detection–non-detection and inventory data further linked scores to observed frequencies under their respective survey designs for fish and trees, but covered only these two groups. Moreover, calibration did not remove observation-effort bias from the nationwide stack. NANAKUSA therefore adjusts stacked scores for observation effort and displays them as within-region percentile ranks of relative biodiversity potential, separately from observed records, rather than as expected species richness, absolute occupancy probabilities or confirmed inventories.

The need for validated biodiversity information extends beyond public conservation planning. As nature-related disclosure under the TNFD framework and science-based target setting under SBTN become more widely adopted, companies and other non-state actors are increasingly expected to link their activities to specific locations and to prioritize areas for action (TNFD, 2023; SBTN, 2024). However, spatially explicit biodiversity information whose predictive reliability and limitations have been evaluated remains limited (TNFD, 2025; Mandle et al., 2024). Externally validated and regularly updated prediction systems of the kind described here can supply such information for arbitrary user-defined areas, provided that their outputs are treated as screening information for designing baseline surveys and continued monitoring, rather than as substitutes for causal assessments of impacts or measurements of restoration outcomes. If observations collected through such surveys are shared, non-state actors could become not only users of biodiversity data but also contributors who help fill local observation gaps. The current implementation focuses on Japan, but the same design principles could be applied wherever open observation records, environmental data and independent validation datasets are available.

This study has several important limitations. First, for the taxonomic groups modelled using shared representations, alien and native species were included in the same training process. Environmental representations learned from well-observed alien species may have supported predictions for data-poor native species. However, because alien species were concentrated in urban areas, they may also have introduced signals associated with urban environments into the shared representations. Comparing models trained with and without alien species, and evaluating their effects on predictions for native species, particularly data-poor species, is therefore an important direction for future research.

Second, most of the training data consisted of presence-only observations. Biases in observation effort and imperfect detection could not be fully removed through the design of background sites or correction for observation effort. External validation also did not cover all 8,297 deployed species, and predictive performance varied substantially among data-poor species added from the taxonomic long tail. Predictions for these species should therefore be treated as candidates requiring field verification, rather than as definitive distribution information. In addition, internal calibration values represent observed frequencies within evaluation datasets composed of presence records and background sites, rather than absolute occupancy probabilities in nature. Calibration using external datasets that included confirmed absences or explicit detection processes was also limited to fishes and trees. Third, the output resolution of the prediction surfaces should be distinguished from their effective ecological resolution. Predictions were produced on a 20 m grid, but the native resolutions of the input covariates ranged from approximately 10 m to 1 km. The 20 m output should therefore not be interpreted as evidence that the models can directly distinguish habitat differences at that spatial scale. Fourth, the external validation of survey recommendations differed in both scale and metric from their implementation in the public platform. The independent evaluation used an unadjusted data-poor-species potential aggregated across nationwide 0.1° cells, whereas the web platform displays an approximately 500 m local recommendation layer after removing the effect of observation effort. Prospective field validation is therefore needed to determine whether the published recommendation layer can efficiently detect data-poor species and improve predictions after model retraining. Implementing a monthly cycle of observation, prediction and renewed survey is a strength of this study, but its long-term effects should be evaluated using the observation records and model-update histories that accumulate over time.

Overall, this study shows that large-scale citizen-science observations can be transformed into a cyclical biodiversity information infrastructure that integrates distribution modelling, external validation, survey recommendations and continued model updating. Its value lies not in providing a single highly accurate map, but in establishing a nationwide feedback system that repeatedly updates predictions and enables the iterative refinement of observation, validation and decision-making while making the basis and limitations of those predictions explicit. The platform could serve as an information infrastructure connecting citizens, public agencies and companies in support of nature-positive action. Where sufficient biodiversity observations, environmental information and independent validation datasets with different observation processes are available, the same design principles could be applied beyond Japan.

## 4. Methods

### 4.1. Biodiversity observation data and spatial framework

This study covered terrestrial and coastal areas throughout Japan. Species occurrence records were collected from the Global Biodiversity Information Facility (GBIF), iNaturalist, eBird, the Ministry of the Environment’s Ikimono Log, and field surveys conducted by researchers. For each record, the scientific name, observation date, geographic coordinates and data source were standardized, followed by taxonomic harmonization and quality control. Records of humans and captive animals originated almost entirely from iNaturalist casual-grade observations that were ingested into the version-1 observation master. The observation master has since been rebuilt to exclude casual-grade records, and subsequent model versions are trained on this casual-free master. After duplicate removal, the canonical data store contained 2,322,578 records from 37,995 species across the full period. Of these, 1,461,834 records from 26,869 species within the 2020–2025 observation window were used for model training and evaluation. Species were assigned to seven taxonomic groups: mammals, birds, reptiles and amphibians, fishes, arthropods, other invertebrates, and plants and fungi.

For the main model training, we initially selected 5,608 species with at least 20 occurrence records during the observation window. The inclusion of species with fewer records and the selection of species for final deployment are described in the next section. Record counts, species numbers, study periods, species-selection criteria and numbers of deployed species by data source and taxonomic group are provided in Supplementary Table 1. Taxonomic harmonization, duplicate handling, coordinate quality control and species-selection procedures are described in Supplementary Methods 1.

We used a nominal set of 41 environmental covariates to explain species distributions. These covariates represented climate, topography, land use, distances to features such as rivers, railways and coastlines, soil, and satellite-derived vegetation indices. Because of data-joining and coverage limitations, approximately 29 covariates contributed effectively to the analyses. Each covariate was aligned to the nationwide prediction grid according to the spatial resolution of the available source data. The native resolutions of the input datasets ranged from approximately 10 m to 1 km. The names, data sources, native resolutions and temporal coverage for each covariate are provided in Supplementary Table 2; preprocessing procedures are summarized in Supplementary Methods 1.

A regular grid with 20 m cells was used to display nationwide predictions and to aggregate outputs within user-defined regions. However, 20 m represents only the unit of prediction output and display, and not the effective spatial resolution of the input covariates or model accuracy. A 0.1° grid was used to evaluate nationwide observation effort and survey-prioritization strategies, whereas an approximately 500 m spatial unit was used to display local survey recommendations. External evaluations used survey sites, locations or spatial units that matched the design of each independent dataset. The construction of these spatial units and the aggregation procedures across resolutions are described in Supplementary Methods 1.

### 4.2. Species distribution models and species selected for deployment

For each taxonomic group, we developed a shared-representation deep-learning model that predicted many species simultaneously (Chen et al., 2017; Brun et al., 2024). Environmental covariates at each location were passed through shared layers, and the model produced an occurrence score for each species within the taxonomic group using a multilabel classification framework (Hu et al., 2025). This design allowed the model to learn environmental responses shared across species and use them in species-level predictions.

The training data consisted of locations with recorded species occurrences and background locations without occurrence records (Phillips and Elith, 2013; Guillera-Arroita et al., 2015). Therefore, non-occurrences in the training matrix did not represent confirmed absences from field surveys. Instead, they represented pseudo-absences in a presence-background design that contrasted occurrence records with background locations. Raw model outputs were therefore treated as relative habitat-suitability scores compared with background locations, rather than as absolute occurrence or occupancy probabilities (Phillips and Elith, 2013; Guillera-Arroita et al., 2015). Background sampling, spatial partitioning, model architecture, loss functions, training iterations, hyperparameters and model-selection procedures are described in Supplementary Methods 1 and Supplementary Table 3.

We first trained models for seven taxonomic groups using 5,608 species with at least 20 occurrence records in Japan during 2020–2025. For fishes, arthropods, and plants and fungi, we then applied long-tail pooling by adding species with fewer occurrence records to the shared training set. For these groups, the nominal minimum number of records was reduced from 20 to 5. After data quality control and model-input requirements were applied, the minimum number of records among species actually included in training was eight for arthropods and for plants and fungi. The deployed fish roster retains species with fewer records within the 2020–2025 window; the effective minimum per taxonomic group is given in Supplementary Table 1.

Based on taxon-specific performance evaluations (Fig. 2a), we selected shared-representation deep models for birds, fishes, arthropods, other invertebrates, and plants and fungi, and species-specific ensemble models for mammals and reptiles and amphibians. The final nationwide deployment included 8,297 species. Of these, 7,987 species were predicted using shared-representation deep models for five taxonomic groups, and 173 species were predicted using species-specific ensemble models for two groups. The remaining 137 species accounted for the difference between the taxon-level total of 8,160 species and the overall deployment total of 8,297 species. These comprised 129 species handled by rare-species fallback models using species-specific ensembles, including 20 birds, 39 arthropods, 50 other invertebrates, and 20 plants and fungi, together with eight species retained in an earlier arthropod model configuration. This composition matched the final deployment manifest, in which mammals and reptiles and amphibians used species-specific ensembles and the other five taxonomic groups used shared-representation deep models.

Nationwide prediction surfaces were generated for all 8,297 deployed species, and all were made available through the web application. Supplementary Data 1 provides the scientific name, taxonomic group, number of occurrence records, number of observed cells, model route, long-tail-pooling status, internal-evaluation status and deployment status for each species. Taxon-specific selection criteria, selected model families and final numbers of deployed species are provided in Supplementary Table 1.

Species were not filtered or separated by native status during model training, evaluation or species-level deployment. As a consequence of this no-filtering policy, the deployed roster includes *Homo sapiens*, domesticated and zoo-kept taxa, alien species and marine mammals wherever their records met the inclusion thresholds; the full manifest of the 8,297 deployed species is provided in Supplementary Data 1. Native and alien species were modelled together within each taxonomic group, including in shared-representation models where applicable. After deployment, alien species were identified using GRIIS Japan (version 25 March 2026; DOI: 10.15468/nt2yla), with 69 species reassigned as native following manual review because they represented either species native to Japan with introduced populations elsewhere within the country or apparent misclassifications of Japanese native species. Alien species were retained in the species-level predictions and in the biodiversity layer of the web application, which is served as three layers: all species, native species only and alien species only. The conservation-priority layer is computed from native species only; including alien species would leave its ranking almost unchanged (Supplementary Methods 7.4).

### 4.3. Model comparison and evaluation of predictive performance

To evaluate predictive performance in unsampled regions, occurrence and background locations were divided into geographic blocks, and the training, validation and test datasets were separated spatially (Roberts et al., 2017). The same spatial partitions were used for all model comparisons. The main evaluation metric was species-level ROC-AUC, which measured whether the model ranked occurrence locations above background locations for each species (Hanley and McNeil, 1982).

We compared the shared-representation deep model with species-specific random forests trained independently for each species. To assess the robustness of this comparison, we conducted additional analyses using different random seeds and alternative species-specific model configurations. We also performed a fully controlled comparison for 28 species, using identical occurrence records, background locations, environmental covariates and spatial partitions for MaxEnt (Phillips et al., 2006), random forest (Breiman, 2001), graph neural network (Scarselli et al., 2009) and non-spatial multilayer perceptron models (Rumelhart et al., 1986).

To distinguish the effect of shared representation from that of the spatial graph, we conducted ablation analyses comparing shared and species-specific models, graph neural networks and multilayer perceptrons, alternative graph-connectivity definitions, and different numbers of species trained jointly (Meyes et al., 2019). Details of the model comparisons, robustness analyses, controlled 28-species comparison and ablation analyses are provided in Supplementary Methods 2.

To assess how strongly non-spatial random splitting inflated performance estimates, we compared species-level ROC-AUC under spatial-block and non-spatial random partitions while holding the species set, environmental covariates, background locations and model settings constant (Roberts et al., 2017). Only species that could be evaluated under both partitioning schemes were included in the paired comparison.

We evaluated the effects of long-tail pooling by comparing models trained only on species with at least 20 occurrence records with models that also included species with fewer records. For well-observed focal species, we tested whether adding data-poor species changed predictive performance. For species with few records, we evaluated how many species became newly evaluable and measured their ROC-AUC. This design allowed us to distinguish the effect of long-tail pooling on the performance of already modelled species from its effect on expanding the range of species that could be predicted. Training conditions and evaluation procedures for long-tail pooling are described in Supplementary Methods 2.

Statistical comparisons among models were based on paired species-level ROC-AUC values for the same species. Details of the statistical analyses, including effect sizes, confidence intervals, P values, multiple-testing corrections and equivalence tests used to assess whether differences were small, are provided in Supplementary Methods 2 and Supplementary Table 4.

To evaluate model generalization across observation designs and detection processes, we used species inventories from certified natural sites, ANEMONE MiFish environmental DNA surveys, structured surveys from Monitoring Sites 1000, complete forest inventories and the National Forest Inventory as external datasets. Site-level community AUC measured whether species recorded at a site were ranked near the top of the candidate-species list for the same taxonomic group. Species-level AUC measured whether sites where a species was recorded or detected were ranked above sites where it was not recorded, not detected or considered absent. Because these metrics represent different use cases, we did not interpret them as directly comparable measures of the same type of performance.

For certified natural sites, we conducted a sensitivity analysis to assess possible inflation caused by spatial overlap between training and evaluation data. Site–species evaluation pairs were excluded when a training record of the same species occurred within 2 km of the site boundary (Roberts et al., 2017). For birds, we also retrained the models after removing training records of the same species from the surroundings of evaluation sites, allowing us to assess the effect of information leakage from nearby records.

For ANEMONE MiFish, repeated water-sampling detection histories were used to estimate occupancy states while accounting for the detection process, and the resulting evaluation was compared with one based on simple detection and non-detection. For the complete forest inventories and the National Forest Inventory, recorded and non-recorded sites were defined according to the design of each survey. The survey design, study period, numbers of sites and species, taxonomic and spatial matching, evaluation criteria and treatment of non-records for each external dataset are described in Supplementary Methods 3 and Supplementary Table 5.

### 4.4. Calibration of prediction scores and the stacked biodiversity index

In addition to evaluating the ranking performance of species-level scores, we examined the correspondence between score magnitude and observed frequency in the evaluation data. For nationwide deployment, scores from occurrence and background locations were pooled by taxonomic group and model route, and calibration functions were fitted using isotonic regression (Niculescu-Mizil and Caruana, 2005; Phillips and Elith, 2010). Calibration performance was evaluated using expected calibration error (ECE) and the Brier score (Brier, 1950), and was compared before and after calibration (Guo et al., 2017; Konowalik and Nosol, 2021; Hui et al., 2023).

The geographic transferability of the calibration relationship was evaluated separately from the nationwide deployment using mean ensembles of random forest and logistic regression models for four taxonomic groups in central Japan. Calibration functions were fitted using the western third of the evaluation area and evaluated in the eastern third, with the central third excluded as a buffer zone. For the ANEMONE MiFish environmental DNA survey and the complete forest inventories, additional calibration functions were estimated using detection–non-detection and presence–absence data, respectively, to match the observation design of each dataset. The preparation of calibration data, geographic partitioning, buffer zones, isotonic regression, ECE, Brier score and external-data calibration procedures are described in Supplementary Methods 4. Calibration results by taxonomic group and external dataset are provided in Supplementary Table 6.

Calibrated scores should not be interpreted as true occupancy probabilities at individual locations or as absolute occurrence probabilities in nature. Instead, they represent calibrated relative scores corresponding to observed frequencies under the observation design used in each evaluation, and were used to compare and stack model outputs across locations.

For the stacked layers, seven human and domesticated taxa (*Homo sapiens*, *Bos taurus*, *Capra hircus*, *Equus caballus*, *Felis catus*, *Gallus gallus* and *Oryctolagus cuniculus*) were removed from the 8,297 deployed species. Although these taxa contributed only 1.92% of the national stacked total, they dominated the highest observation-effort-adjusted residuals: *Homo sapiens* alone accounted for 88.8% of the stacked value in the top-ranked residual cell. *Sus scrofa* was retained as the native Japanese wild boar. The 542 species classified as alien were then also excluded. Nine marine mammals (*Enhydra lutris*, *Eumetopias jubatus*, *Megaptera novaeangliae*, *Orcinus orca*, *Phoca largha*, *Phoca vitulina*, *Sagmatias obliquidens*, *Tursiops aduncus* and *Tursiops truncatus*) were additionally removed because the prediction nodes carry a land mask but no habitat mask and these species therefore received scores in inland cells. They contributed 0.35% of the national stacked total but dominated individual residual cells, including the second-and third-ranked cells. For each prediction cell, calibrated scores for the remaining 7,739 native species were summed to calculate the stacked biodiversity prediction and observation-effort-adjusted residuals reported here. In the web application, the biodiversity layer is served as three layers (all 8,290 species, native species only and alien species only); alien species account for 7.97% of the national all-species stacked total (Supplementary Methods 5). The resulting stacked value represents relative biodiversity potential predicted by the models, rather than absolute expected species richness or a sum of occupancy probabilities.

Japan was divided into 0.1° grid cells, and the stacked calibrated scores and the number of existing occurrence records within each cell were log-transformed. The stacked prediction was regressed against the number of occurrence records to estimate the value expected from observation effort in each cell. The difference between the observed stacked value and this expectation was calculated as a residual, allowing the relative level of biodiversity potential to be compared among areas with similar observation effort. Procedures for aggregation to 0.1° cells, selection of eligible cells, log transformation, regression and residual calculation are described in Supplementary Methods 5.

In the web application, this observation-effort-adjusted index was rescaled within each region specified by the user. The displayed values therefore represent relative rankings within the selected region and do not indicate absolute species richness. Values calculated for different user-defined regions cannot be compared directly on a common absolute scale.

### 4.5. Evaluation of survey-prioritization strategies

We evaluated survey-prioritization strategies from two perspectives: their ability to improve overall model performance and their ability to efficiently detect species with few observation records. To evaluate model improvement, we first withheld part of the training records and retrained the models. Candidate sites were then ranked using different strategies, and the withheld records were returned to the training data in order of site priority. This retraining experiment compared three strategies: a composite score combining prediction uncertainty, gaps in observation effort and data-poor-species potential; prediction uncertainty alone; and random selection. Selection based on the existing distribution of observations and data-poor-species potential alone was examined separately in the species-discovery analysis described below. At each step, the models were retrained, and we measured how rapidly species-level ROC-AUC on a fixed test dataset recovered towards its value before records were withheld.

To evaluate the efficiency of detecting data-poor species, we used independent detection records from ANEMONE MiFish and Monitoring Sites 1000 that were not used for model training. We focused on 227 species with fewer than 20 occurrence records in the training data. Nationwide 0.1° grid cells were ranked using unadjusted data-poor-species potential, prediction uncertainty, the composite score, the existing distribution of observation sites or random selection. Assuming that cells were surveyed in rank order, we compared the cumulative number of data-poor species detected and the number of cells required to include a given proportion of the target species. The site-selection strategies, sequential retraining procedure and evaluation of cumulative discovery using external records are described in Supplementary Methods 6.

The survey-priority index implemented in the web application was calculated by summing calibrated prediction scores across deployed data-poor species and regressing this value against existing observation effort. The survey-priority score for each cell was defined as the residual, representing the extent to which data-poor-species potential exceeded the level expected from similar observation effort. High values therefore indicate areas where predicted signals for data-poor species are concentrated beyond what can be explained by existing observation effort. The external evaluation directly tested the unadjusted data-poor-species potential. The discovery efficiency of the residualized score implemented in the web application was not directly evaluated under the same external-validation design.

The discovery efficiency of the survey-prioritization strategies was evaluated using nationwide 0.1° grid cells. In the web application, an approximately 500 m local recommendation surface based on the same principle was also displayed to support the selection of field-survey locations. However, discovery efficiency at this approximately 500 m display unit was not directly validated by the evaluation conducted at the 0.1° scale. Definitions of the survey-priority scores, spatial units and procedures used to transform the nationwide surface into the local surface are provided in Supplementary Methods 6.

### 4.6. Implementation in the web application

The web application allowed users to define a target region by selecting a prefecture, municipality or registered site, or by drawing an arbitrary polygon on the map. For each selected region, the application displayed existing observation records, a candidate-species list based on predicted scores, species-level prediction maps, a biodiversity prediction map, a conservation-priority map and a survey-priority map.

Species-level prediction maps and candidate-species lists included all deployed species. In contrast, the biodiversity prediction map was calculated by stacking calibrated scores across the 8,290 species described above and adjusting the resulting values for observation effort; it is served as all-species, native-only and alien-only layers. The conservation-priority map summed the calibrated scores of native species only, after weighting each species by the inverse of its nationwide mean calibrated score. The survey-priority map used the residualized data-poor-species potential after accounting for observation effort.

These regional indices were rescaled within each user-defined target region. They were therefore intended for comparing candidate locations within a region and could not be used directly for absolute comparisons of biodiversity among different regions. The conservation-priority map was not a comprehensive conservation prioritization that directly incorporated extinction risk, future land-conversion risk, conservation costs or management feasibility.

Existing observation records and model predictions were displayed separately to prevent predicted candidate species from being interpreted as a confirmed regional inventory. Species-level predictions on the 20 m grid and survey recommendations at an approximately 500 m unit were outputs intended to identify relative candidate locations within a region. They did not guarantee ecological accuracy at their respective display resolutions. Procedures for defining target regions, calculating each output, setting spatial resolutions, normalizing values within regions and applying display constraints are described in Supplementary Methods 7 and Supplementary Table 7.

### 4.7. Use of generative artificial intelligence

Generative AI tools, including Claude Code (Anthropic) and Codex (OpenAI), were used as assistive tools during software and web-platform development, code generation and debugging, data-processing and analytical workflow development, documentation, and English translation and editing of the manuscript. All AI-generated or AI-modified code, analytical procedures and written material were reviewed, tested and verified by the authors before use. Decisions concerning study design, model specification, validation, statistical interpretation and scientific conclusions were made by the authors, who take full responsibility for the accuracy, integrity and reproducibility of the work. Generative AI tools were not treated as sources of research data or scientific evidence.

## Data availability

The processed data supporting the findings of this study are provided with this paper as Source Data and Supplementary Data 1. These files include species-level model-performance metrics, external-validation and calibration results, aggregated indices for 0.1° grid cells, survey-prioritization results and the data underlying all figures. Species-level model-performance metrics are provided where retained. Some ablation analyses are available only as run-level summaries, and the staged-retraining experiment is available as seed-averaged recovery curves, as specified in the Source Data README. Source data are provided with this paper. Occurrence records used for model development were obtained from GBIF, iNaturalist, eBird, the Ministry of the Environment of Japan’s Ikimono Log, and surveys conducted by the authors and collaborating researchers. Publicly available records can be accessed from the original data providers. Data sources, observation periods and record counts are provided in Supplementary Table 1; publicly available records can be accessed from the original data providers. Exact occurrence coordinates for threatened or otherwise sensitive species are not publicly released because disclosure could increase the risk of poaching, illegal collection or habitat disturbance. These restricted data are available from the corresponding authors for legitimate research or conservation purposes, subject to the terms of the original data providers, an assessment of risks to sensitive species and completion of a data-use agreement prohibiting the redistribution or disclosure of sensitive locations. To support independent verification and reuse, we also provide free programmatic access to the deployed models upon request. Approved applicants can query species-level predictions; regional biodiversity, conservation-priority and survey-priority indices; and observation-effort layers for any location or user-defined area. This access enables applicants to reproduce the platform outputs reported here and conduct independent validation. The same access mechanism is used to provide large prediction outputs and accompanying documentation that are not deposited in a public repository because they contain species-sensitive information. Applications are reviewed to verify the applicant’s identity and confirm that the proposed use is legitimate and is not intended to recover protected locations of threatened species. Access is granted under terms prohibiting the redistribution or disclosure of sensitive locations. Requests should be submitted through https://mapry.co.jp/contact/.

## Code availability

Reference implementations of the two model families used in this study, together with a synthetic dataset on which they can be run, are available on Zenodo at 10.5281/zenodo.22062792 under the Creative Commons Attribution 4.0 International License. The implementations cover model fitting and are provided to allow the specifications described in Supplementary Methods 1 to be inspected and executed. Because the implementations are trained on synthetic data, they do not reproduce the performance values reported here. The species distribution models were fitted using the production codebase of the biodiversity platform operated by Mapry Inc. This codebase is not publicly available because it forms part of the operational software of a public service and implements the aggregation and coarsening rules used to protect the withheld locations of threatened species, as described under Data availability. This study introduces no new algorithm. The model architecture, loss function, optimizer settings, training schedule, background-sampling procedure, spatial-partitioning scheme and taxon-specific deployment rules are described in Methods and Supplementary Methods 1–7. The processed data underlying the reported results and programmatic access to the deployed models are available as described under Data availability.

## Acknowledgements

We thank the many citizen scientists across Japan whose sustained efforts to observe and document biodiversity over many years made this study possible.

## Author contributions

K.Y. collected the data, developed the web platform, conducted all analyses and prepared all figures. Y.F. drafted the manuscript and led the writing process. Y.F, K.U. and M.K.H. contributed to discussions that shaped the direction of the study and critically revised the manuscript. All authors reviewed and approved the final manuscript.

