## Supplementary Information for "Closing the biodiversity observation-to-action loop"

**A national biodiversity platform linking citizen observations to conservation planning and future surveys**

### Supplementary Methods 1

Observation data, environmental covariates, model construction and deployment

#### SM1.1. Biological occurrence data

We collected species occurrence records covering terrestrial and coastal areas throughout Japan from the Global Biodiversity Information Facility (GBIF), iNaturalist, eBird, the Ministry of the Environment’s Ikimono Log, and field surveys conducted by researchers. After duplicate removal, the canonical data store contained 2,322,578 records from 37,995 species across the full period. We defined 2020–2025 as the observation window for the analyses; 1,461,834 records from 26,869 species within this window were used for model training and evaluation. These two sets of values refer to different data populations and should not be cited interchangeably. Record and species counts by data source and taxonomic group are provided in Supplementary Table 1. Full dataset citations, versions, access dates and identifiers are provided in the supplementary data-source documentation.

For each record, the scientific name, observation date, latitude, longitude and data source were standardized to a common format. We excluded records for which the observation date or geographic coordinates could not be interpreted, records located outside Japan, and records for which the taxonomic name could not be resolved reliably to species level. Records of the same species with matching observation dates and spatial locations were cross-checked across data sources, and duplicate observations were merged into a single record.

#### SM1.2. Taxonomic standardization

Scientific names were standardized against the GBIF Backbone Taxonomy. Taxonomic authorship, spelling variants and infraspecific ranks were reconciled during this process. Records identified only by a Japanese common name were converted to scientific names using a curated correspondence table. Names that could not be assigned unambiguously to a single species were excluded from model training.

Standardized species were assigned to seven taxonomic groups: mammals, birds, reptiles and amphibians, fishes, arthropods, other invertebrates, and plants and fungi. These groups formed the basic units for model training, deployment and evaluation.

Species of conservation concern, including threatened species, were retained for model training, but their precise occurrence locations were not disclosed directly through the web application. The coordinate-handling procedures used for public release are described in Supplementary Methods 7.

#### SM1.3. Spatial framework

We used a regular grid of 20 × 20 m cells to generate nationwide predictions and aggregate outputs within arbitrary user-defined regions. Occurrence locations, background locations, environmental covariates and model predictions were assigned to the same 20 m grid, and each cell was identified by its centre coordinates.

The 20 m grid was generated using the appropriate zone of the Japan Plane Rectangular Coordinate System under JGD2011. Each location was projected into the plane rectangular coordinate system and assigned to a grid at 20 m intervals, after which the cell centre was transformed back to geographic coordinates. The 20 m grid represents the unit used for prediction output, display and regional aggregation; it does not represent the effective spatial resolution of the environmental covariates or the ecological accuracy of the models.

A 0.1° grid was used for nationwide stacked predictions, correction for observation effort and evaluation of survey-prioritization strategies. An approximately 500 m spatial unit was used for local survey recommendations in the web application. External validation used sites, locations or survey plots corresponding to the survey design of each dataset.

#### SM1.4. Environmental covariates

Each prediction cell was assigned a nominal set of 41 environmental covariate columns. These comprised seven climate variables, three elevation variables, two slope variables, 11 forest-resource variables, five land-use composition variables, including one mean vegetation-naturalness variable, three distances to land-use classes, five distances to other geographic features, four soil variables and one satellite-derived vegetation index. The model input dimensionality during training and deployment was therefore nominally 41. The names, definitions, units, data sources, native resolutions and temporal coverage of these variables are provided in Supplementary Table 2; preprocessing, missing-value imputation and standardization are summarized below.

The nominal and effective numbers of covariates differed. The 11 forest-related variables, including forest resources, tree height, stand age, stem density and growing-stock volume, were included in the input matrix. However, these data were joined by exact matching of coordinates to six decimal places and were not snapped to the 20 m grid. Consequently, the proportion of non-missing values at training nodes was only 0.03–0.34%. Missing values were imputed with a constant value of zero, and the permutation importance of these variables remained at the noise level. These variables therefore contributed effectively no information to the models. All performance values reported in this study were consequently obtained without effective forest-structure covariates.

Similarly, the distance-to-river variable had a non-missing proportion of 0.4671 among the 150,000 training nodes used for the deployed core bird model. All nodes were missing in 29 prefectures for which the prefecture-level source data could not be obtained; the corresponding proportion was 0.9945 in the corrected version of the layer. The mean vegetation-naturalness variable was calculated without excluding sentinel values and therefore cannot be interpreted as a valid index of vegetation naturalness. After excluding these ineffective or uninterpretable variables, approximately 29 covariates materially supported the reported results: 12 climate and terrain variables, four land-use composition variables, eight distance variables, four soil variables and one vegetation index.

For environmental layers with a native resolution coarser than 20 m, each 20 m cell was assigned the value of the source mesh or raster cell in which it occurred. Land-use composition variables were calculated by summarizing land-use classes within a spatial neighbourhood around each prediction cell. Distance variables were calculated as the nearest distance from the cell centre to the relevant land-use class or geographic feature.

Missing values were imputed using either a representative value or zero, depending on the properties of the variable. Continuous variables were standardized using means and standard deviations estimated from the training data, and the same transformations were applied to the validation, test and nationwide prediction data.

#### SM1.5. Selection of training species and long-tail pooling

For the initial model training, we selected 5,608 species with at least 20 occurrence records in Japan during the 2020–2025 observation window. The initial number of training species in each taxonomic group is provided in Supplementary Table 1.

For fishes, arthropods, and plants and fungi, we applied long-tail pooling by adding species with fewer occurrence records to the shared training set. For these three groups, the nominal minimum number of records required for inclusion was reduced from 20 to 5.

The initial number of training species, the number added through long-tail pooling, the final number of deployed species and the selected model family for each taxonomic group are provided in Supplementary Table 1. Supplementary Data 1 provides the scientific name, taxonomic group, number of occurrence records, number of observed cells, long-tail-pooling status and final model route for every deployed species.

#### SM1.6. Shared-representation deep models

For each of the seven taxonomic groups, we trained a shared-representation deep model that simultaneously predicted many species within the group. The input was the nominally 41-dimensional environmental covariate vector described in SM1.4, of which approximately 29 dimensions contributed effectively to the predictions.

The shared trunk comprised fully connected layers with 512, 512, 256 and 128 units. Each layer was followed by batch normalization, a rectified linear unit activation and dropout. Residual connections, with linear projections where required to match dimensions, were introduced between adjacent layers. The dropout rate was 0.3. A single output layer transformed the final 128-dimensional shared representation into one logit for each training species within the taxonomic group.

The target matrix was a multilabel matrix spanning all training species in the taxonomic group. An entry was assigned a value of 1 when the focal species had been recorded in a cell and 0 otherwise. A value of 0 did not represent an absence confirmed by a structured survey; it represented a pseudo-absence within the presence–background design. The sigmoid outputs of the model were therefore treated as relative scores conditional on the occurrence and background sampling design, rather than as absolute probabilities of occurrence or occupancy in nature.

#### SM1.7. Background sampling and spatial holdout

The training data consisted of cells with recorded species occurrences and background cells. For the shared-representation models, background cells were generated in the vicinity of occurrence locations. Random spatial jitter was added to the latitude and longitude of each occurrence location, and the resulting locations were assigned to the 20 m grid. Cells that coincided with occurrence cells and duplicate background cells were removed.

As a general rule, the number of background cells was twice the number of occurrence cells. The number of occurrence cells was capped at 4,000 per species, and the total number of training cells was capped at 150,000 per taxonomic group.

Occurrence and background cells were partitioned into ten spatial blocks using K-means clustering based on their geographic coordinates. Entire blocks were assigned to the training, validation or test set, thereby reducing the likelihood that nearby locations were split among different datasets (Roberts et al., 2017).

#### SM1.8. Model training

The shared-representation deep models were trained using a binary cross-entropy loss that accounted for imbalance among species and between occurrence and background cells. We used the AdamW optimizer (Loshchilov and Hutter, 2019) with an initial learning rate of 0.002, a weight decay of 0.0001, a batch size of 16,384 and a maximum of 300 epochs. Training was stopped when performance on the validation data failed to improve for a specified period, and the model weights from the epoch with the best validation performance were retained.

The model architecture is described in SM1.6, and the loss function, optimization settings and early-stopping procedure are specified in this section.

#### SM1.9. Per-species ensemble models

Based on taxon-specific performance evaluations, we selected per-species ensemble models, rather than shared-representation deep models, for nationwide deployment of mammals and reptiles and amphibians. These models were trained independently for each species.

The ensemble comprised three base learners: a spatial graph attention network (GAT; hidden dimension of 128, four attention heads and weighted focal loss), a random forest with 60 trees, a maximum depth of 10 and class_weight="balanced", and logistic regression with max_iter=600 and class_weight="balanced" (Breiman, 2001; Lin et al., 2017; Veličković et al., 2018). Their predictions were combined using a fixed weighted average rather than a simple mean:

0.5 × graph attention network + 0.25 × random forest + 0.25 × logistic regression.

A habitat mask was then applied to the weighted prediction to obtain the final score. The full ensemble was used only for species with at least eight occurrence nodes. For species with fewer occurrence nodes, or for which one or more base learners failed to train, predictions from the graph attention network alone were used.

An equal-weight average of random-forest and logistic-regression predictions was used in a separate pathway to fit isotonic calibrators for reptiles and amphibians and for the rare-species fallback models. It was not the method used to combine predictions in the deployed ensemble. This model route was consistent with the final deployment manifest: shared-representation deep models were used for five taxonomic groups, whereas per-species ensembles were used for mammals and reptiles and amphibians. The base learners, hyperparameters and ensemble settings are specified in this section. Background-cell sampling is described in SM1.7.

#### SM1.10. Deployment and national prediction

Based on taxon-specific performance evaluations, we selected shared-representation deep models for birds, fishes, arthropods, other invertebrates, and plants and fungi. Per-species ensemble models were selected for mammals and reptiles and amphibians.

Nationwide predictions were generated for 8,297 species. Of these, 7,987 species were predicted using shared-representation deep models for five taxonomic groups, and 173 species were predicted using per-species ensemble models for two groups. The remaining 137 species accounted for the difference between the taxon-level total of 8,160 species and the overall deployment total of 8,297 species. These comprised 129 species handled by rare-species fallback models using per-species ensembles—20 birds, 39 arthropods, 50 other invertebrates, and 20 plants and fungi—and eight species retained in an earlier arthropod model configuration. The 8,297 deployed species corresponded to the species made available through the web application.

Nationwide prediction surfaces for each species were produced on the 20 m grid. Because the raw outputs of both the shared-representation deep models and the per-species ensemble models depended on the presence–background design, they were not interpreted as absolute probabilities of occurrence. Instead, they were passed to a subsequent calibration procedure described in Supplementary Methods 4.

The initial number of training species, the number added through long-tail pooling, the final number of deployed species and the selected model family for each taxonomic group are provided in Supplementary Table 1. The shared-model architecture and training settings are described in SM1.6 and SM1.8, respectively; the composition of the per-species ensembles is described in SM1.9; and background-cell sampling and spatial partitioning are described in SM1.7. Taxon-specific deployment routes are summarized in Supplementary Table 3. The final model route for each of the 8,297 deployed species is provided in Supplementary Data 1.

### Supplementary Methods 2

Model comparisons, long-tail pooling and ablation analyses

#### SM2.1. Comparison of shared-representation and per-species models

We compared the discrimination performance of the shared-representation deep models with that of species-specific random forests trained independently for each species. Both model types were evaluated using the same spatial holdout defined in Supplementary Methods 1. The primary evaluation metric was species-level ROC-AUC, which measures the ability of a model to rank occurrence cells above background cells. Species were considered evaluable when the test data contained at least three occurrence cells and three background cells. AUC values obtained for the same species were paired and compared within each taxonomic group. Both model types were trained and evaluated on the same 41-column covariate matrix, standardized with statistics fitted on the training nodes only, and on the same spatial partition; the asymmetric predictor sets described in SM2.2 apply only to the pipeline-level comparison. The species-specific baseline was a random forest with 80 trees, a maximum depth of 12 and class_weight="balanced", fitted independently for each species; the per-species ensembles described in SM1.9 are the models selected for nationwide deployment and were not used as this baseline.

To assess the robustness of the comparison, we retrained the models using different random seeds and performed additional comparisons using alternative configurations of the species-specific models. We also compared AUC values obtained under spatial-block and non-spatial random partitioning while holding the species set, environmental covariates, background cells and model settings constant. This comparison was restricted to species that were evaluable under both partitioning schemes.

For the robustness analysis, models were retrained using two random seeds (seeds 0 and 1). The seed controlled both the resampling of background cells and the spatial-block partitioning. Occurrence cells were identical between the two seeds, and the resulting spatial partitions converged on essentially the same geographic blocks. Comparisons between the shared-representation model and the species-specific random forest were summarized separately for each seed, and we confirmed that the direction of the taxon-specific conclusions was consistent between seeds. AUC values were averaged across seeds for each species only in the internal cross-validation of the deployed models. As alternative species-specific configurations for birds, we additionally considered the better-performing species-specific model selected from a random forest tuned to 300 trees and a histogram-based gradient-boosting model.

#### SM2.2. Pipeline-level and controlled model comparisons

For the pipeline-level comparison, we used an independent complete forest inventory comprising 125 tree species. Prediction scores generated by the shared-representation model, MaxEnt (Phillips et al., 2006) and species-specific random forests were evaluated at the same external survey plots. For each species, we calculated ROC-AUC to quantify discrimination between plots at which the species was recorded and those at which it was not recorded. Because each model used its own training data, background locations and environmental covariates, this analysis compared the complete deployment pipelines rather than model architectures alone. In particular, the environmental predictor sets were asymmetric. To make nationwide species-specific retraining computationally feasible, MaxEnt and the species-specific random forests used an approximately 17-dimensional subset comprising climate variables, the five highest-ranked distance variables, soil variables and the vegetation index. The deployed shared-representation model used the nominal set of 41 input columns, of which approximately 29 contributed effectively.

To compare model architectures under controlled conditions, we selected the four species with the largest numbers of records from each of the seven taxonomic groups, giving a total of 28 species. For each species, we used identical occurrence records, background cells, environmental covariates comprising the nominal 41 columns (approximately 29 effective columns), and spatial partitions to compare a graph neural network, a non-spatial multilayer perceptron, a random forest and MaxEnt. Training and evaluation data were separated at the level of spatial blocks. The deep models were evaluated inductively: neither the nodes nor the edges in the held-out evaluation blocks were used during training. The number of species included and a summary of the comparison results are provided in Supplementary Table 4.

#### SM2.3. Long-tail pooling

We evaluated the effects of long-tail pooling for fishes, arthropods, and plants and fungi. Species with at least 20 occurrence records were defined as focal species, whereas those with 5–19 records were defined as low-record species. We compared a condition in which only the focal species were trained jointly with a condition in which the low-record species were added to the shared training set. The cells, environmental covariates, spatial partitions and model architecture were held constant between the two conditions; only the set of species included in joint training was changed.

For the focal species, species-level ROC-AUC values before and after the addition of low-record species were paired to assess whether predictive performance was retained. For the external evaluation using the complete tree inventory for plants and fungi, cell-level predictions from the pre-pooling model were reconstructed from the retained nationwide prediction surfaces and isotonic calibrators using the same procedure as the original evaluation code. The reconstructed values closely reproduced the reported species-level AUC values (Pearson’s *r* = 0.9985; mean absolute error = 0.0043). The before- and after-pooling arms were then paired using the same 59 plots and the calibrated score column in both arms.

For low-record species, we evaluated the number of species that became evaluable under the spatial holdout and calculated their species-level ROC-AUC values. A species was considered evaluable when the test data contained at least three occurrence cells and three background cells. Whether AUC exceeded 0.6 was used only as a post hoc descriptive measure of performance and was not used as a criterion for selecting species for deployment.

Retention of performance among focal species was assessed by pooling paired species-level differences across multiple random seeds and applying two one-sided tests (TOST; Lakens, 2017) with an equivalence margin of ±0.01 AUC. Here, *n* denotes the number of species-by-seed pairs rather than the number of species. Equivalence was supported only for arthropods. It was supported both in the uncontrolled analysis (*n* = 778 pairs, mean Δ = −0.0047, 90% CI [−0.0079, −0.0014], *P* = 0.0036) and in the controlled analysis in which early stopping was based only on the focal species (*n* = 262 pairs, mean Δ = +0.0022, *P* = 0.025). For fishes, equivalence was not supported in either the uncontrolled analysis (*n* = 218 pairs, mean Δ = +0.0070, *P* = 0.25) or the controlled analysis (mean Δ = +0.0039, 90% CI [−0.0025, +0.0104], *P* = 0.061). Equivalence tests were not conducted for the other five taxonomic groups.

#### SM2.4. Ablation analyses

To distinguish the gain from shared representation learning from that attributable to the spatial graph, we conducted ablation analyses using identical cells, the nominal set of 41 environmental covariates (approximately 29 effective covariates), and the same spatial holdout. To assess the effect of shared representation learning, we compared a multi-output shared GNN with independently trained species-specific GNNs, and a multi-output shared MLP with independently trained species-specific MLPs. To assess the contribution of the spatial graph, we compared the shared GNN with a shared MLP that did not perform neighbourhood aggregation over a graph.

We evaluated the effects of graph structure by altering the connectivity of the proximity graph, weighting edges according to distance, adding edges along river corridors, and connecting cells within the same catchment. We also held six focal species constant while increasing the total number of jointly trained species from 6 to 12, 25, 50 and 100, thereby testing how the number of species included in shared training affected predictive performance for the focal species.

These ablation analyses were conducted within a rectangular region covering central Japan and the Kanto region (135.0–141.5° E, 34.0–37.6° N). We used the 25 species with the largest numbers of records in each taxonomic group and repeated the analyses using three random seeds. All deep models were evaluated inductively, such that information from the held-out evaluation blocks was not used for message passing during training.

The shared GNN and shared MLP were trained in separate series of runs. However, the MLP runs used the same node construction and feature-extraction procedures as the GNN runs and shared the study extent, species list, random seed, background-cell generation and spatial blocks, which were derived deterministically from the seed. For reptiles and amphibians, the AUC values for the common random-forest baseline stored by the two series were exactly identical for 256 of 285 evaluation units, with a maximum difference of 9.5 × 10⁻⁵. This row-wise agreement confirmed that the two models had been evaluated on the same units, and we therefore conducted a paired comparison between the shared GNN and shared MLP for this taxonomic group only. For birds, the common baseline values did not agree (median difference = 0.018) because differences in the maximum number of nodes produced non-identical evaluation units; we therefore did not conduct a paired comparison for birds.

The two series of runs were also not matched in the procedure used to select hyperparameters, which involved grid search for the GNN but default values for the MLP; the number of training epochs, which was 50 for the GNN and 120 for the MLP; or the data used to fit feature standardization, which were restricted to the within-fold training data for the GNN but included all nodes for the MLP. The difference in standardization would be expected to favour the shared MLP. Irrespective of its direction, these configuration differences mean that the higher performance of the shared GNN reported in the main text (paired median difference = +0.016) cannot be attributed specifically to the spatial graph.

Proximity graphs were constructed using taxon-specific connection radii ranging from 100 to 300 m: 200 m for mammals, 300 m for birds, 150 m for reptiles and amphibians, and 100 m for fishes. In the original graphs, for which background cells were generated using spatial jitter of ±0.009° (approximately 1 km), 22% of bird nodes, 53% of mammal nodes and 59% of reptile and amphibian nodes were isolated. We therefore repeated the analyses for mammals and reptiles and amphibians using connectivity-repaired graphs generated by reducing the jitter to 0.001° (approximately 110 m), which reduced the proportion of isolated nodes to approximately 2%. The main conclusion that neighbourhood aggregation over the spatial graph provided no clear gain was unchanged. The connectivity-repaired bird graph was too dense to train successfully. For mammals, the multi-species graph model did not differ significantly from the species-specific graph model either before or after connectivity repair (P = 0.107 and P = 0.491, respectively; Supplementary Table 4(d)), and after repair its difference from the random-forest baseline was not significantly different from zero (−0.001, *P* = 0.68).

#### SM2.5. Statistical analyses

Model comparisons were based on paired ROC-AUC values obtained for the same species. Taxon-specific performance was summarized using the median species-level AUC, and effect sizes between models were calculated as the median of the paired species-level differences. Differences were tested using the Wilcoxon signed-rank test. We calculated 95% confidence intervals by bootstrap resampling with species as the resampling unit. When ablation analyses included multiple random seeds, repeated values from the same species were treated as belonging to the same cluster. For the ablation contrasts quoted in the running text and in SM2.4, the effect estimate is the median of the per-species mean differences, so that the estimate, the confidence interval and the test all refer to the species as the unit. The effect estimates and confidence intervals reported in Supplementary Table 4(d) instead summarise all paired records, with the Wilcoxon test computed on the per-species means.

For the taxon-specific model comparisons, the seven paired Wilcoxon tests, one for each taxonomic group, were treated as a single family and adjusted using the Holm method. The differences remained significant after adjustment for birds (adjusted *P* = 0.003) and arthropods (adjusted *P* < 0.001). The results for fishes (adjusted *P* = 0.077) and other invertebrates (adjusted *P* = 0.086) were significant only before adjustment. The adjusted values were calculated retrospectively from the stored test results; the analysis outputs contained only the unadjusted *P* values. Comparisons against multiple baselines in the validation of survey recommendations were adjusted using the Bonferroni method in accordance with the preregistered analysis, whereas no multiple-testing adjustment was applied to the ablation analyses.

The number of evaluable species, effect sizes, 95% confidence intervals, *P* values, multiplicity-adjusted *P* values and equivalence-test results for each analysis are provided in Supplementary Table 4. Species- and replicate-level values are provided in the Source Data where retained. For analyses available only as run-level summaries or seed-averaged results, the level of data aggregation is specified in the Source Data README.

### Supplementary Methods 3

External validation using independent biodiversity datasets

#### SM3.1. General evaluation framework

We evaluated whether the nationally deployed models could provide effective rankings when applied to datasets with observation designs and detection processes different from those of the training data. The evaluation used certified natural sites, the ANEMONE MiFish environmental DNA survey, Monitoring Sites 1000, complete forest inventories and the National Forest Inventory. The survey period, spatial unit, number of sites, number of species, criteria for evaluability and interpretation of non-records for each dataset are provided in Supplementary Table 5. The deployment-matched prediction artefacts used for external validation covered 8,292 of the 8,297 deployed species.

Taxonomic names in the external datasets were standardized using the same taxonomic framework as in SM1.2. Only species whose scientific names matched those in the nationwide deployment were included in the evaluation. For sites defined by polygons, species-level scores were aggregated across the prediction cells within each site. For sites defined by points or survey plots, scores from the corresponding prediction cells were used.

Site-level community ROC-AUC measured whether species recorded at a site were ranked above non-recorded deployed species from the same taxonomic group. Species-level ROC-AUC measured whether sites at which a species was recorded or detected were ranked above sites at which it was not recorded, not detected or considered absent. AUC was calculated only when both positive and negative observations were available. Because these two AUC measures represent different use cases, they were not compared directly as equivalent measures of performance.

#### SM3.2. Certified natural sites

We obtained site-level species inventories from the certification materials for the certified natural sites and matched scientific and Japanese common names to the nationally deployed species. Species-level scores were aggregated across prediction cells whose centres fell within each site-boundary polygon to generate a ranked candidate-species list for each site. If a polygon contained no prediction-cell centre, the prediction from the nearest cell was used.

To examine whether spatial overlap between training records and evaluation sites inflated apparent performance, we excluded site–species pairs for which a training record of the same species occurred within 2 km of the site boundary. We then recalculated both site-level community AUC and species-level AUC. This procedure removed 977 of 4,880 presence records (20.0%). The dataset contained 393 sites and 1,302 species for which species-level AUC could potentially be calculated. Site-level community AUC was calculable for 348 sites before exclusion and 335 sites after exclusion, whereas species-level AUC was calculable for 1,302 species before exclusion and 1,224 species after exclusion.

This leakage-control procedure was partial for three reasons. First, it removed presence records from the evaluation data but did not modify the pseudo-absence component. Second, because the buffer was specified in geographic degrees, its effective east–west width ranged from 1.46 to 1.64 km rather than 2 km. Third, 73.8% of the predictions were represented by the nearest prediction cell outside the site polygon. As an additional analysis for birds, we removed training records of the same species within 2 km of each evaluation site, retrained the models and compared their AUC values with those from models that retained the nearby records.

#### SM3.3. ANEMONE MiFish environmental DNA survey

For the ANEMONE MiFish environmental DNA survey, detection histories from repeated water samples at each site were matched to the nationally deployed species. In the primary evaluation, a species was treated as present at a site if it was detected in at least one replicate and as not detected if it was absent from all replicates. Site-level community AUC and species-level AUC were then calculated using these detection–non-detection labels.

To examine imperfect detection across repeated samples, we fitted a multispecies occupancy model to the detection histories. For the subset of species and sites to which the occupancy model could be applied, species-level AUC was nevertheless calculated using the same detection–non-detection labels as in the primary evaluation. Because non-detection in environmental DNA data does not constitute confirmed absence, the primary evaluation and the occupancy-model subset were treated as distinct evaluations.

Occupancy was estimated using the Bayesian multispecies occupancy model implemented in occumb (Fukaya and Hasebe, 2025). Control samples were excluded, and replicates were restricted to those collected during the same visit. The model was fitted to all 255 fish species. Occupancy (ψ) and sequence relative dominance (φ) were modelled with species-specific intercepts, and sequence capture probability (θ) as a function of the log-transformed, standardized volume of filtered water. We retained the package default priors: a normal prior with mean 0 and precision 1×10⁻⁴ for community-level means and effects shared across species, a uniform prior U(0, 10⁴) for the standard deviations of species-specific effects, and a uniform prior U(−1, 1) for their correlation coefficients. This fit was run in R 4.5.3 with occumb 1.3.0, using JAGS 4.3.2 as the MCMC engine via jagsUI, with four chains of 6,000 iterations each (including 2,000 burn-in iterations) after 1,000 adaptation iterations, thinned by 3, retaining 5,332 posterior samples. Twenty-four species that had not converged in this fit were subsequently re-fitted as a separate subset and the two sets of estimates were merged; the sampler settings of that subset re-fit were not retained. For the 231 species from the documented fit, R-hat was below 1.1 for every species (median 1.007, maximum 1.094). Across the merged set of 255 species, R-hat was below 1.2 for every species and below 1.1 for 252, with a maximum of 1.16; the three species exceeding 1.1 all came from the subset re-fit. Because occumb shrinks species-specific effects towards a community-level distribution estimated from the species included in a fit, estimates from the two fits are not directly comparable, and statistics based on ψ are therefore reported for the documented fit only. The occupancy model was built from species detected at three or more sites, a detection requiring at least ten sequence reads in a replicate; the evaluated subset comprised the 126 of these species for which a nationally deployed model was available. Occupancy was fitted across 137 sites, and species-level AUC was evaluated at the 123 sites for which model predictions were available. Species-level AUC remained based on detection–non-detection labels, with a site classified as positive when the species was detected in at least one replicate; estimated occupancy, ψ, was not used to calculate AUC.

Of the 126 evaluated species, 110 have ψ from the documented fit and 16 from the subset re-fit. Across those 110 species the median ψ was 0.092, and estimated occupancy exceeded the raw detection rate, computed on the same 123-site evaluation frame, for 78 of them. We used ψ as the denominator when calculating PR-gain, defined as PR-AUC divided by prevalence. This produced a more conservative metric than using the raw detection rate as the denominator. Because the raw detection rate was obtained in a separate summary based on a different set of sites, it was not presented as if it represented the same population as ψ. The interpretation of metrics normalized by estimated occupancy and their distributions among species are also being examined in a separate manuscript in preparation.

#### SM3.4. Monitoring Sites 1000

Structured survey records from Monitoring Sites 1000 were matched to the nationally deployed models using scientific names and survey locations. These records did not overlap temporally with the observation window used for model training and were therefore not included in the training data. Species recorded at each site were treated as present, whereas non-recorded deployed species from the same taxonomic group were treated as non-recorded. We then calculated site-level community AUC and species-level AUC. Because non-records could reflect imperfect detection, they were not interpreted as confirmed absences.

#### SM3.5. Complete forest inventories

For the complete forest inventories, tree species recorded within a survey plot were treated as present, whereas target species not recorded in the complete inventory of that plot were treated as absent. Using species-level scores from the prediction cell corresponding to each plot, we calculated species-level ROC-AUC to evaluate whether plots at which each tree species was present were ranked above plots at which it was absent.

#### SM3.6. National Forest Inventory

For the National Forest Inventory, tree-species records from standardized forest survey plots were matched to the nationally deployed species. For each tree species, plots at which the species was recorded were treated as positive observations, whereas eligible survey plots at which it was not recorded were treated as non-records. Species-level ROC-AUC was then calculated.

Because the National Forest Inventory and complete forest inventories differed in survey design and in the certainty with which non-records could be interpreted as absences, their AUC values were presented as separate external evaluations. Further details of the survey design, the procedure used to map Japanese common names to accepted scientific names, the distribution of species-level AUC values and the attribution of failure modes among species with low AUC values are being reported in a separate manuscript in preparation.

### Supplementary Methods 4

Calibration of species-level prediction scores

#### SM4.1. Spatially separated calibration data

In addition to evaluating the ranking performance of the models, we assessed the correspondence between raw output scores and observed frequencies in the evaluation data. Calibration used presence records and background locations from spatial holdout data that had not been used for model training. Geographic transferability was evaluated by dividing the evaluation locations at the 33rd and 67th percentiles of longitude. The calibration function was fitted using the western third, the central third was excluded as a buffer, and performance was evaluated in the eastern third.

This analysis was a separate experiment restricted to a rectangular region in central Japan (135.0–141.5° E, 34.0–37.6° N). It included the 25 species with the largest numbers of records in each taxonomic group, subject to a minimum of 15 records. The scores being calibrated were generated by a mean ensemble of random forest and logistic regression models rather than by the shared-representation deep models. The experiment was conducted only for mammals, birds, reptiles and amphibians, and plants and fungi. This longitudinal partitioning was specific to the calibration experiment and was not used to calibrate the nationally deployed models. The experiment was not conducted for fishes, arthropods or other invertebrates.

Metadata describing the geographic partition were retained in the analysis artefacts for only three groups: birds, reptiles and amphibians, and plants and fungi. The mammal results were obtained from an earlier output that did not contain this metadata. Consequently, although the boundaries and buffer were defined by the 33rd and 67th percentiles of evaluation-point longitude as described above, the retained artefacts do not allow us to confirm that the mammal results were generated using the identical three-part partition.

#### SM4.2. Isotonic calibration and evaluation metrics

For each taxonomic group, we fitted a monotonic isotonic regression function (Niculescu-Mizil and Caruana, 2005) using the raw species-level output scores and their corresponding presence–background labels. The fitted function was applied to the evaluation data in the eastern partition to transform raw scores into calibrated scores. When an evaluation score fell outside the range of scores in the calibration data, it was assigned the value at the nearest endpoint of the fitted calibration function. Because isotonic regression is monotonic, it largely preserved the ranking of model outputs while mapping their magnitudes to observed frequencies.

Calibration performance was evaluated using expected calibration error (ECE) and the Brier score, and both metrics were compared before and after calibration. To calculate ECE, prediction scores were divided into bins, and the absolute difference between the mean prediction and observed frequency in each bin was aggregated with weights proportional to the number of observations in that bin. The Brier score was calculated as the mean squared difference between the predicted score and the binary label. Calibration results before and after transformation are provided by taxonomic group in Supplementary Table 6, and calibration curves are shown in Supplementary Fig. 5.

For the spatially separated calibration experiment, ECE was calculated using ten equal-width bins spanning [0, 1]. External calibration was evaluated using equal-frequency bins defined by quantiles of the predicted scores: ten nominal bins for the environmental DNA data and eight nominal bins for the complete forest inventories. Duplicate quantile boundaries were merged, so the effective number of bins could be smaller than the nominal number. Because both the binning procedure and the number of bins differed, ECE values cannot be compared directly between the spatially separated calibration experiment and the external-calibration evaluations, or between the environmental DNA and complete-forest-inventory evaluations.

#### SM4.3. Calibration using external presence–absence data

The labels in the internal holdout data were based on a presence–background design comprising occurrence records and background locations. Consequently, calibrated scores derived from these data did not directly represent absolute probabilities of occurrence in nature. We therefore fitted additional isotonic calibration functions using detection–non-detection data from the ANEMONE MiFish environmental DNA survey for fishes and presence–absence data from the complete forest inventories for plants. These functions matched the observation design of the corresponding external dataset. ECE and Brier scores before and after external calibration are provided in Supplementary Table 6.

For both external datasets, sites were divided into western and eastern subsets at the median longitude. Calibration functions were fitted and evaluated in both directions, from west to east and from east to west. Calibration was applied only when ECE improved in both directions; otherwise, it was not used. The calibration function ultimately applied to the deployment artefacts was refitted using all cells pooled together after passing this bidirectional validation. Thus, separation between training and evaluation data was used to assess generalization, whereas the final deployed calibrator was fitted to the complete external dataset.

The resulting quantities remained specific to the corresponding survey design. For fishes, the calibrated value represented the probability of environmental DNA detection at the approximately 10-km grid-cell scale under the ANEMONE survey design. For plants, it represented the probability of presence within a complete-inventory plot. Both were retained as separate columns rather than replacing the existing calibrated scores.

#### SM4.4. Interpretation and downstream use

Calibrated scores were interpreted as relative scores corresponding to observed frequencies under the observation design used for calibration. The calibration procedure used for national deployment did not include a geographic split. For each taxonomic group, pairs of raw scores and presence–background labels were pooled across all deployed species and all training cells to fit a single isotonic calibrator. These calibrators were applied to all species in the five taxonomic groups for which shared-representation deep models provided the primary output: birds, fishes, arthropods, other invertebrates, and plants and fungi.

Mammals were excluded from this calibration pathway after their deployment model was changed to the per-species ensemble. The 61 deployed species without a raw-score column in the artefacts corresponded exactly to all deployed mammal species. Separate ensemble-based calibrators were applied to reptiles and amphibians and to the rare-species fallback models, covering 1,163 species in total.

Before-and-after ECE and Brier scores were available only for the four taxonomic groups included in the separate calibration experiment: mammals, birds, reptiles and amphibians, and plants and fungi. These metrics were not obtained for fishes, arthropods or other invertebrates. The deployment manifest recorded the number of calibration applications by taxonomic group only for fishes. For birds, arthropods, and plants and fungi, however, the deployed values reproduced the corresponding taxon-specific calibrators, confirming that calibration had been applied even though application counts were not recorded in the artefacts.

Calibrated scores were not interpreted as the true occupancy probability at a location or as an absolute probability of occurrence in nature. They were used to compare model outputs among locations and to stack predictions across species. The procedures used to stack species-level predictions and correct for observation effort are described in Supplementary Methods 5.

### Supplementary Methods 5

Stacked biodiversity predictions and correction for observation effort

#### SM5.1. Stacked biodiversity predictions

For each prediction cell, we summed the calibrated scores of the deployed native species to calculate a stacked biodiversity prediction. We excluded seven human, domesticated or captive-derived species (*Homo sapiens*, *Bos taurus*, *Capra hircus*, *Equus caballus*, *Felis catus*, *Gallus gallus* and *Oryctolagus cuniculus*) and 542 alien species. We additionally excluded nine marine mammals (*Enhydra lutris*, *Eumetopias jubatus*, *Megaptera novaeangliae*, *Orcinus orca*, *Phoca largha*, *Phoca vitulina*, *Sagmatias obliquidens*, *Tursiops aduncus* and *Tursiops truncatus*), leaving 7,739 native species in the analysis. Wild boar (*Sus scrofa*) was retained as a native wild species. Calibrated species-level scores were obtained using the procedures described in Supplementary Methods 4. The stacked value was treated as a relative index of the extent to which predicted signals from many species overlapped at a location. It was not interpreted as an absolute expected number of species, a sum of occupancy probabilities or a count of confirmed species.

Species were not selected or excluded according to native status during model training or species-level deployment. By contrast, the stacked analysis reported in this section and in Fig. 4 was restricted to native species and additionally excluded the nine marine mammals because the prediction nodes carried a land mask but no habitat mask, causing these species to receive scores in inland cells. The nine species contributed 0.35% of the national stacked total but dominated individual residual cells, including the second- and third-ranked cells. Marine fishes were not removed because primary species-level habitat information distinguishing marine, brackish and freshwater taxa was unavailable. Their contribution therefore remains in the stacked prediction and should be considered when interpreting coastal cells.

The main biodiversity layer in the web application included 8,290 species after removing only the seven human and domesticated or captive-derived species, and was provided as three separate layers: all species, native species only and alien species only. Of these 8,290 species, 542 were classified as alien on the basis of establishmentMeans = Alien in GRIIS Japan. These species accounted for 7.97% of the all-species stacked prediction. We treated 69 species classified as alien by GRIIS as native after manual review; these comprised 27 species native to Japan but introduced into regions outside their native range within the country and 42 Japanese native species that appeared to have been misclassified. Alien species accounted for 7.10% of the training records and had approximately 3.5 times as many records per species as native species.

For the nationwide comparison, calibrated scores at the 20 m prediction nodes were summed directly within each 0.1° cell. The calculations did not pass through an intermediate coarse grid, and no representative value, such as a maximum, was selected for each species within a cell. The aggregation operator was the sum, rather than the mean or maximum.

#### SM5.2. Observation effort and analysis cells

Observation effort was defined as the total number of existing occurrence records from 2020–2025 within each 0.1° cell, using the records from all taxonomic groups integrated as described in SM1. We included 3,971 cells that contained at least five occurrence records and for which a stacked biodiversity prediction was available.

#### SM5.3. Regression and residualization

The stacked biodiversity prediction and the number of occurrence records were each log-transformed. We then fitted a simple linear regression across the nationwide cells, using the log-transformed stacked prediction as the response variable and the log-transformed number of occurrence records as the explanatory variable. For each cell, the residual was calculated as the difference between the observed log-transformed stacked prediction and the value predicted by the regression model. A positive residual indicated that the stacked prediction was higher than expected for a region with a similar level of observation effort, whereas a negative residual indicated that it was lower than expected.

These residuals were calculated to diagnose the relationship between observation effort and the stacked prediction across 7,739 native species after additionally excluding nine marine mammals. They differed from the survey-priority score calculated specifically for data-poor species in terms of the species pool, regression model and intended use (Supplementary Methods 6).

#### SM5.4. Spatial comparison and interpretation

We mapped the uncorrected stacked predictions and the residuals after adjustment for observation effort across Japan and compared the spatial distributions of areas with high values. This analysis assessed the extent to which spatial variation in observation effort associated with citizen-science data remained in a regional indicator obtained by stacking predictions across many species. We did not interpret the uncorrected stacked predictions directly as differences in biodiversity among regions. Likewise, the residuals were not interpreted as an absolute biodiversity index from which all sources of bias other than observation effort had been removed.

The stacked prediction, number of occurrence records, regression-predicted value and residual for each 0.1° cell will be included in the Source Data for Fig. 4b and Fig. 4d. Regional normalization and display within the web application are described in Supplementary Methods 7.

### Supplementary Methods 6

Evaluation and construction of survey recommendations

#### SM6.1. Survey-site selection strategies

We evaluated survey-prioritization strategies from two perspectives: their ability to improve average discrimination across the models and their ability to facilitate efficient discovery of species with few occurrence records. Candidate locations were defined as nationwide 0.1° cells. We considered predictive uncertainty, low observation effort, data-poor-species potential, a composite score combining these quantities, selection following the existing spatial distribution of observation sites, and random selection.

The sets of strategies compared differed between the two evaluations. The staged retraining experiment described in SM6.2 compared three strategies: the composite score, predictive uncertainty and random selection. The evaluation of data-poor-species discovery described in SM6.3 compared five strategies: data-poor-species potential alone, the composite score, predictive uncertainty, selection following the existing distribution of observation sites, and random selection.

Predictive uncertainty was calculated by averaging the binary entropy of species-level predictions across species within each cell. Low observation effort was ranked such that cells containing fewer existing occurrence records received higher priority. Data-poor-species potential was calculated by stacking the calibrated prediction scores of species with few occurrence records within each cell. The composite score combined the ranks of predictive uncertainty, low observation effort and data-poor-species potential.

#### SM6.2. Staged retraining experiment

To evaluate improvement in overall model performance, we removed a subset of the existing training records and retrained the models. Candidate cells were scored while these records were withheld, and the withheld records were then returned to the training data in the order determined by each prioritization strategy. At each stage, the models were retrained using the same settings, and species-level ROC-AUC was calculated on a fixed test dataset. Using the ROC-AUC of the model trained before record removal as the reference, we compared how rapidly each strategy restored discrimination performance.

The retraining experiment included fishes, arthropods, and plants and fungi and was restricted to training species with at least 20 occurrence records. The number of nodes was capped at 40,000 per taxonomic group, occurrence cells were capped at 4,000 per species, and background cells were generated using spatial jitter of approximately 1 km. Nodes were divided into training, validation and test data using spatial blocks.

The pool of training nodes was ranked according to each strategy. The models were retrained at five stages using the top 35%, 50%, 65%, 80% and 100% of the ranked nodes. The same training configuration, comprising 120 epochs, was used at every stage, and the median species-level ROC-AUC was calculated on the fixed test data. The experiment was repeated using two random seeds, and the seed-averaged values were used to construct the recovery curves. The three strategies compared in this experiment were descending composite priority score, descending predictive uncertainty and random selection.

#### SM6.3. Independent evaluation of data-poor-species discovery

The efficiency with which data-poor species were discovered was evaluated using detection records from ANEMONE MiFish and Monitoring Sites 1000 that had not been used for model training. We included 227 species that had fewer than 20 occurrence records in the training data and at least one detection record in these external datasets. External detection records were assigned to 0.1° cells, and the candidate cells were ranked according to each strategy.

Assuming that cells were surveyed in rank order, we calculated the cumulative number of target species newly detected among the visited cells. Discovery efficiency was compared using the number of species detected within the top 10% of candidate cells and the number of cells required to include 50% of the target species. The evaluation was based on 78,674 independent detection records across 4,914 candidate cells. Cumulative curves were evaluated at 40 equal budget points, in increments of 122 cells. The exact top decile contained 491 cells; because this point was not included in the saved curves, the reported top-decile readout used 610 cells, the first evaluated budget point at or above 491.

The data-poor-species potential used in this evaluation had not been adjusted for observation effort. Cumulative discovery curves for each strategy are shown in Supplementary Fig. 6, and cell-level rankings and detection data will be provided in the corresponding Source Data.

#### SM6.4. Survey-priority score used in the web application

The survey-priority index implemented in the web application was derived by regressing data-poor-species potential—the stacked calibrated prediction scores of deployed species with few occurrence records—against the number of existing occurrence records in the same cell. The survey-priority score for each cell was defined as the residual, representing the extent to which data-poor-species potential exceeded the value expected from the prevailing level of observation effort. High values therefore indicated areas in which predicted signals for low-record species were concentrated beyond what could be explained by existing observation effort alone.

For the approximately 500 m surface displayed in the web application, the data-poor-species pool comprised 2,678 of the 5,608 training species for which predictions were retained and that had no more than 50 nationwide occurrence records. The nationwide 0.1° surface used a different pool comprising 3,828 species with at least 15 but fewer than 50 records. The species pools used for the two surfaces were therefore not identical.

Data-poor-species potential, $L$, and observation effort, $E$, were transformed as $\log\left( 1+L \right)$ and $\log\left( 1+E \right)$, respectively. We regressed $\log\left( 1+L \right)$ against a third-degree polynomial of $\log\left( 1+E \right)$. The resulting residuals were rank-normalized to obtain the survey-priority score.

The external-data evaluation directly assessed the unadjusted data-poor-species potential. The discovery efficiency of the residualized score deployed in the web application was not evaluated directly under the same external-validation design.

#### SM6.5. National and local recommendation layers

The discovery efficiency of the survey-prioritization strategies was evaluated using nationwide 0.1° cells. To support the selection of field-survey locations, the web application also displayed an approximately 500 m local recommendation surface based on the same general principle. Because the nationwide and local surfaces differed in spatial unit and normalization domain, their numerical values were not compared on a common scale.

The local surface used cells measuring 0.005541° in longitude by 0.004492° in latitude, corresponding to approximately 500 m. Regression against observation effort and rank normalization of the residuals were performed independently within each 1° × 1° tile. Within each cell, data-poor-species potential, (L), was calculated by summing prediction scores across the data-poor-species pool. Each prediction was evaluated at one representative point per cell. For this surface, the scores were sigmoid outputs to which the taxon-specific isotonic calibrators had not been applied.

The version underlying the statistics reported here comprised 108 tiles and 1,116,980 cells. The version currently served through the web application was produced in a separate build using a common nationwide grid; consequently, its numbers of tiles and cells differ from those reported here.

Discovery efficiency at the approximately 500 m display scale was not directly validated by the evaluation conducted using 0.1° cells. Region selection, normalization and display within the web application are described in Supplementary Methods 7.

### Supplementary Methods 7

Web application, regional outputs and interpretation

#### SM7.1. Region selection and spatial aggregation

We implemented the validated and calibrated model outputs in a web application that allowed municipalities, site managers and members of the public to use the predictions without specialist geographic information system software. Users could specify a target region by selecting a prefecture, municipality or registered site, or by drawing an arbitrary polygon on the map. Occurrence records, species-level predictions and regional indicators corresponding to the selected region were extracted and displayed as regional reports and maps.

Grid points were generated at the designated resolution within the bounding box of the selected region. Inclusion within the regional polygon was determined using a point-in-polygon test. Thus, inclusion was based on the location of the cell centre rather than intersection with the region or the proportion of cell area contained within it.

#### SM7.2. Observed records, candidate species and species maps

Existing occurrence records and model predictions were displayed separately for each selected region. Observed species were summarized from records in the occurrence dataset described in SM1 that were located within the selected region. The predicted candidate-species list was generated by aggregating calibrated prediction scores for deployed species within the region and ranking the resulting regional scores. A species-level prediction map on the 20 m grid was displayed for each candidate species. Predicted candidate species were not included in the confirmed regional inventory and were explicitly identified as model-derived candidates.

The regional score for each candidate species was calculated as the sum of its calibrated scores across cells within the selected region, interpreted operationally as the expected number of occupied cells. Neither the maximum nor the mean score was used. Species were included in the candidate list when this sum was at least 2.0; the threshold was reduced to 1.0 for species included on a Red List. On the prediction map for each candidate species, cells with values of at least 50% of the maximum score within the selected region were displayed as locally suitable areas.

For species of conservation concern, precise occurrence locations were not displayed directly but were either spatially coarsened or withheld. Prediction maps were displayed independently of the procedures used to protect observed locations.

#### SM7.3. Biodiversity prediction map

The biodiversity prediction map used a regional indicator obtained by stacking calibrated scores across deployed species and adjusting their relationship with observation effort. The procedures used to calculate the stacked prediction and correct for observation effort are described in Supplementary Methods 5. Within the web application, this indicator was normalized within the user-defined region and displayed as relative variation in biodiversity potential among cells in that region.

Normalization within the selected region used percentile ranks, with tied values assigned the midrank of their tie group. Min–max scaling was not used because the raw scores had a narrow distribution; min–max scaling compressed most values towards the lower end of the scale and resulted in poor visual differentiation.

#### SM7.4. Conservation-priority map

The conservation-priority map was generated by weighting the calibrated prediction score of each species by the inverse of its nationwide mean calibrated score and summing the weighted values within each cell. This procedure assigned relatively high values to cells in which species with low nationwide mean calibrated scores overlapped. The resulting values were displayed after normalization within the selected region.

For species (s), the weight was defined as the inverse of its mean calibrated score across nationwide prediction cells:

wₛ = 1 / (p̄ₛ + 10⁻⁴)

The conservation-priority score for cell (c) was then calculated as

prio(c) = Σₛ wₛ pₛ,꜀

This is an additive benefit function that does not convert prediction scores into binary presence–absence values using a threshold.

For the application-wide display scale, values were normalized by the 99th percentile of the pooled cell-level distribution from a sample of municipalities selected at regular intervals from the nationwide order of municipality codes. The default sample comprised 12 municipalities; owing to the implementation, approximately the final 160 municipalities in the ordered list were not represented in this sample. Normalized values were clipped to [0, 1]. Rank classes were defined using the 95th, 85th, 65th and 35th percentiles of the same reference distribution. Colours on the map were assigned according to percentile ranks within the user-selected region.

We did not report threshold-based community metrics, including predicted species richness, $N_{\tau}$, or the associated rarity-weighted value, $V_{\tau}$. The conventional threshold of 0.5 lay above the maximum range of the calibrated scores, which varied among taxonomic groups from 0.06 to 0.36. The deployed implementation did not classify species as present using an absolute threshold, and threshold constants retained in the source code were not invoked.

The conservation-priority map is a relative indicator of locations where predictions for species with comparatively restricted distributions overlap. It is not a comprehensive measure of conservation priority incorporating extinction risk, future land-conversion risk, conservation cost, land tenure or management feasibility.

The conservation-priority map was calculated using native species only. Including alien species produced virtually no change in its spatial pattern (Spearman’s $\rho=0.9997$ across 0.1° cells; 97% overlap among the top 100 cells). This occurred because the range-rarity weighting assigns greater weight to species with restricted distributions, and alien species did not receive high rankings. The highest-ranked alien species was 1,433rd among all deployed species, and no alien species occurred in the top 1%.

#### SM7.5. Survey-priority map

The survey-priority map used a residualized data-poor-species potential obtained by stacking predictions for species with few occurrence records and adjusting for existing observation effort. The construction of the score and the nationwide 0.1° and approximately 500 m local recommendation surfaces are described in Supplementary Methods 6. Within the web application, the approximately 500 m local surface was normalized within the selected region and displayed as candidate locations for additional surveys.

The external-data evaluation directly assessed the data-poor-species potential before adjustment for observation effort. Neither the deployed residualized score itself nor its discovery efficiency at the approximately 500 m scale was directly validated under the same external-evaluation design.

#### SM7.6. Interpretation and display constraints

The biodiversity prediction, conservation-priority and survey-priority outputs were all normalized within the user-defined region. Their displayed values therefore represent ranks or relative values for comparing candidate cells within that region and cannot be compared among different selected regions on a common absolute scale.

The 20 m species-level predictions represent the units used for display and regional aggregation; they do not guarantee model accuracy at that spatial resolution. Similarly, the approximately 500 m survey-priority surface is a display layer intended to support the selection of field-survey locations. The evaluation results obtained using nationwide 0.1° cells cannot be assumed to apply directly at the approximately 500 m scale.

Candidate-species lists and prediction maps indicate the possible presence of species that have not yet been recorded. They were not interpreted as confirmed occurrence records or as evidence of absence. The definition, inputs, spatial unit, normalization, intended use and main limitations of each output are summarized in Supplementary Table 7.

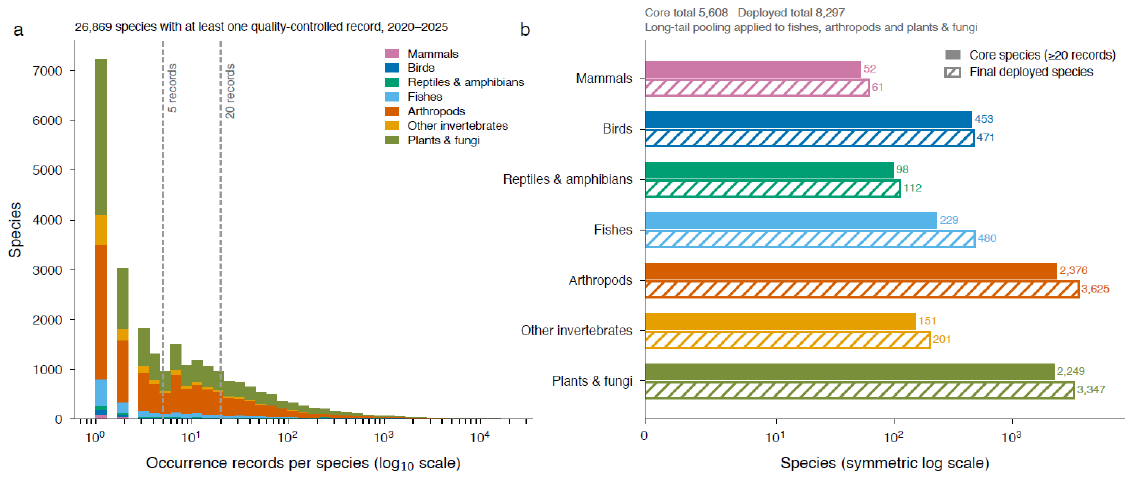

**Supplementary Fig. 1 | Long-tailed occurrence records and expansion of deployed taxonomic coverage.** a, Distribution of quality-controlled occurrence records among 26,869 species observed in 2020–2025, stacked by taxonomic group. Dashed lines indicate 5 and 20 records. b, Numbers of core species with at least 20 records and final deployed species in each group. Long-tail pooling was applied to fishes, arthropods, and plants and fungi, contributing to an increase from 5,608 core species to 8,297 deployed species. The horizontal axis in b uses a symmetric logarithmic scale.

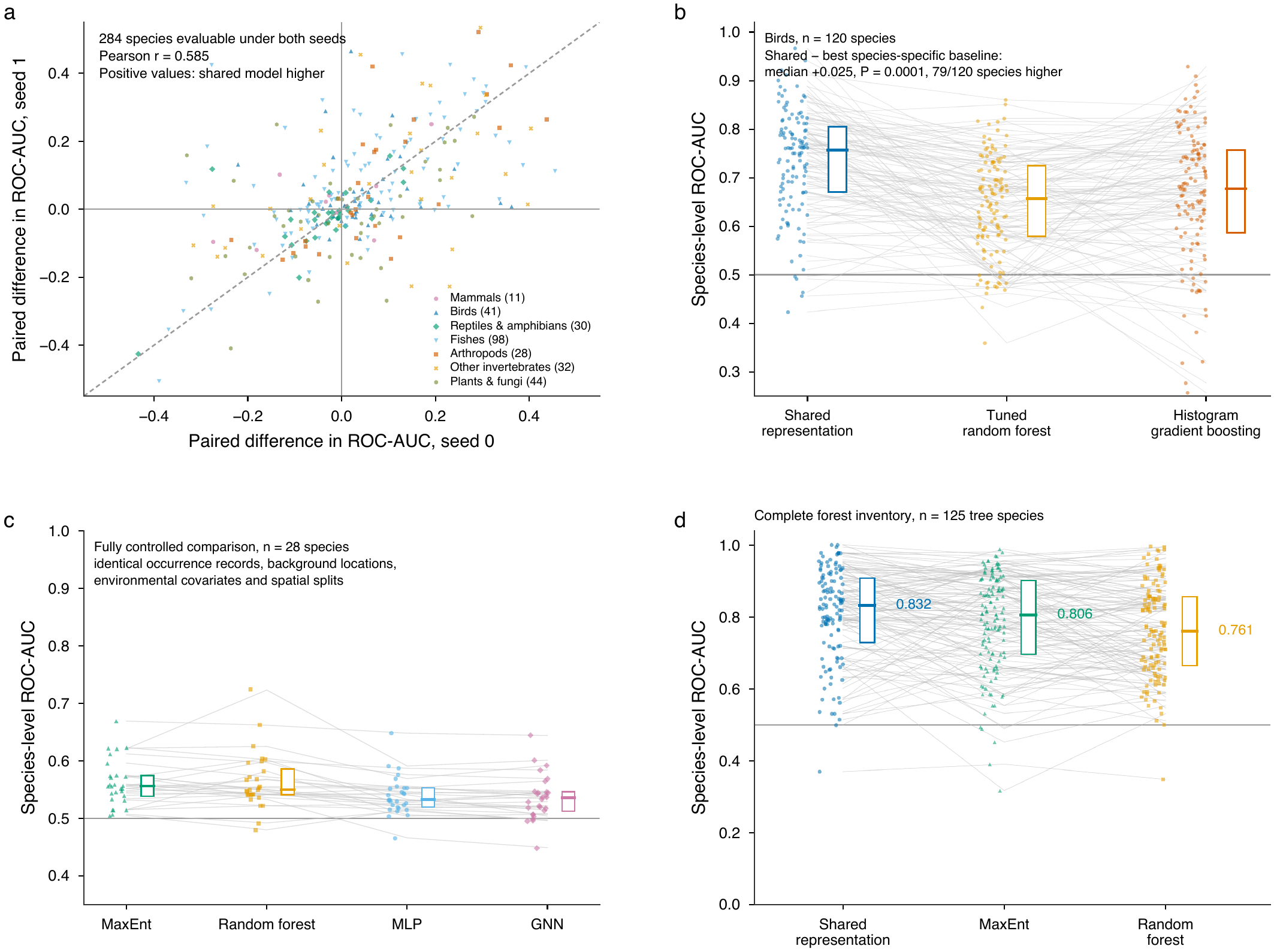

**Supplementary Fig. 2 | Robustness and benchmark comparisons of shared-representation models.** a, Paired differences in species-level ROC-AUC between the shared-representation model and the species-specific random forest under seeds 0 and 1 for 284 evaluable species; positive values favour the shared model and the dashed line denotes equality between seeds. b, Paired comparison for 120 bird species against a tuned random forest and histogram-based gradient boosting. c, Fully controlled comparison of four model families for 28 species using identical records, background cells, covariates, and spatial splits. d, Pipeline-level comparison at complete forest-inventory plots for 125 tree species. Predictor sets and training pipelines differed in d. Points represent species; boxes show the interquartile range and central lines show medians. The horizontal line marks ROC-AUC = 0.5.

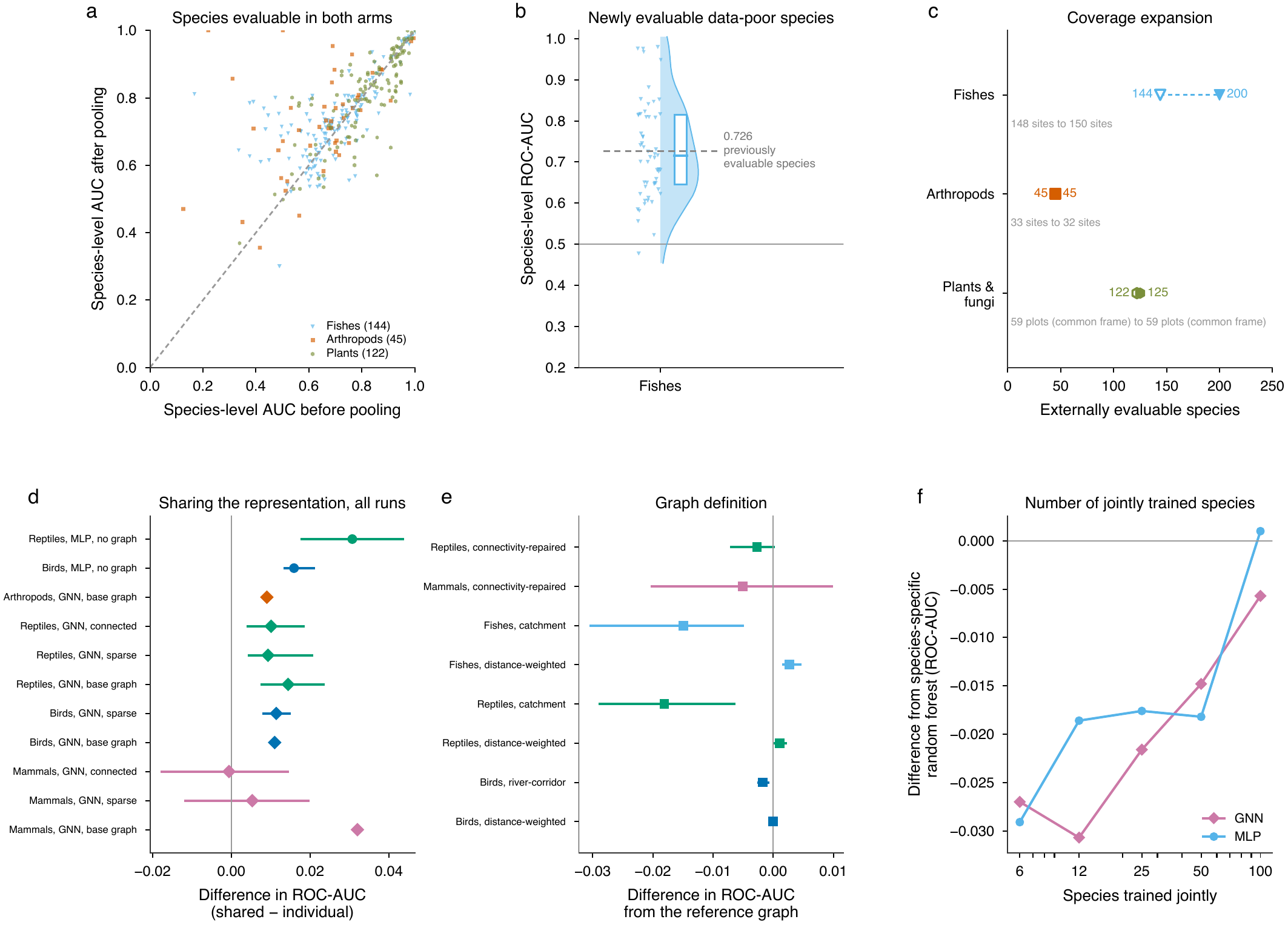

**Supplementary Fig. 3 | Long-tail pooling and ablation analyses.** a, Species-level ROC-AUC before and after pooling for species evaluable in both conditions; the dashed line denotes equality. b, ROC-AUC for 56 fish species that became externally evaluable after pooling; the dashed line shows the median for previously evaluable fish species. c, Numbers of externally evaluable species before and after pooling. d, Differences between shared and species-specific representations across ablation runs; positive values favour sharing. e, Changes relative to each reference graph under alternative graph definitions. f, Difference from the species-specific random forest for six fixed focal species as the number of jointly trained species increased. Points show effect estimates and horizontal bars show 95% confidence intervals where available. Run-specific differences in model configuration are detailed in Supplementary Methods 2.4.

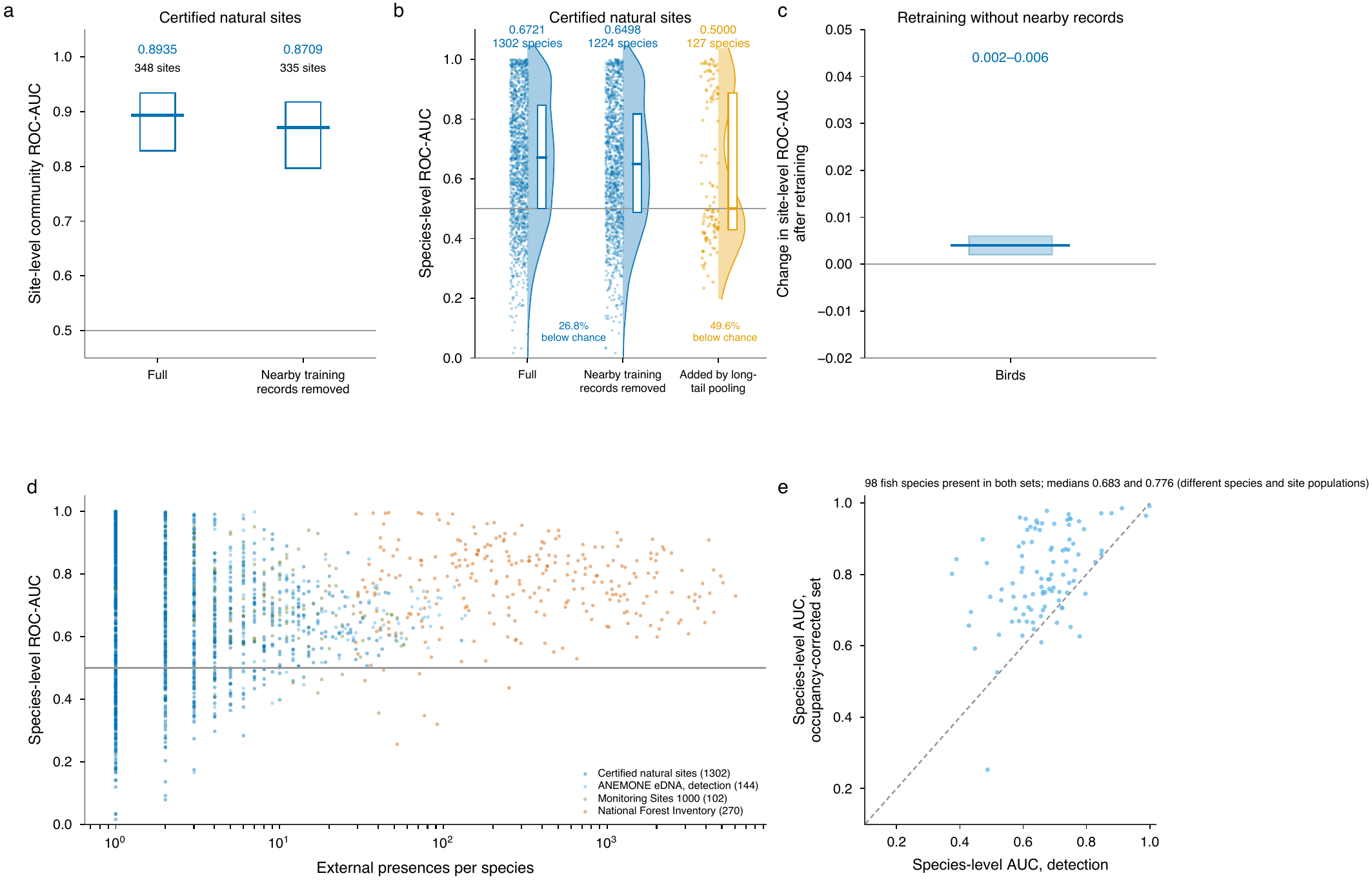

**Supplementary Fig. 4 | External validation and sensitivity to spatial overlap and detection.** a, Site-level community ROC-AUC at certified natural sites in the full and 2-km-controlled evaluations. b, Species-level ROC-AUC for the full and 2-km-controlled certified-site evaluations and for species added by long-tail pooling. c, Change in bird site-level ROC-AUC after retraining without nearby same-species records. d, Species-level ROC-AUC in four external datasets in relation to the number of external presences. e, Detection-based and occupancy-subset ROC-AUC for 98 fish species represented in both ANEMONE evaluations. Non-records and sampling frames differ among datasets; the comparisons are therefore descriptive rather than estimates on a common absolute scale.

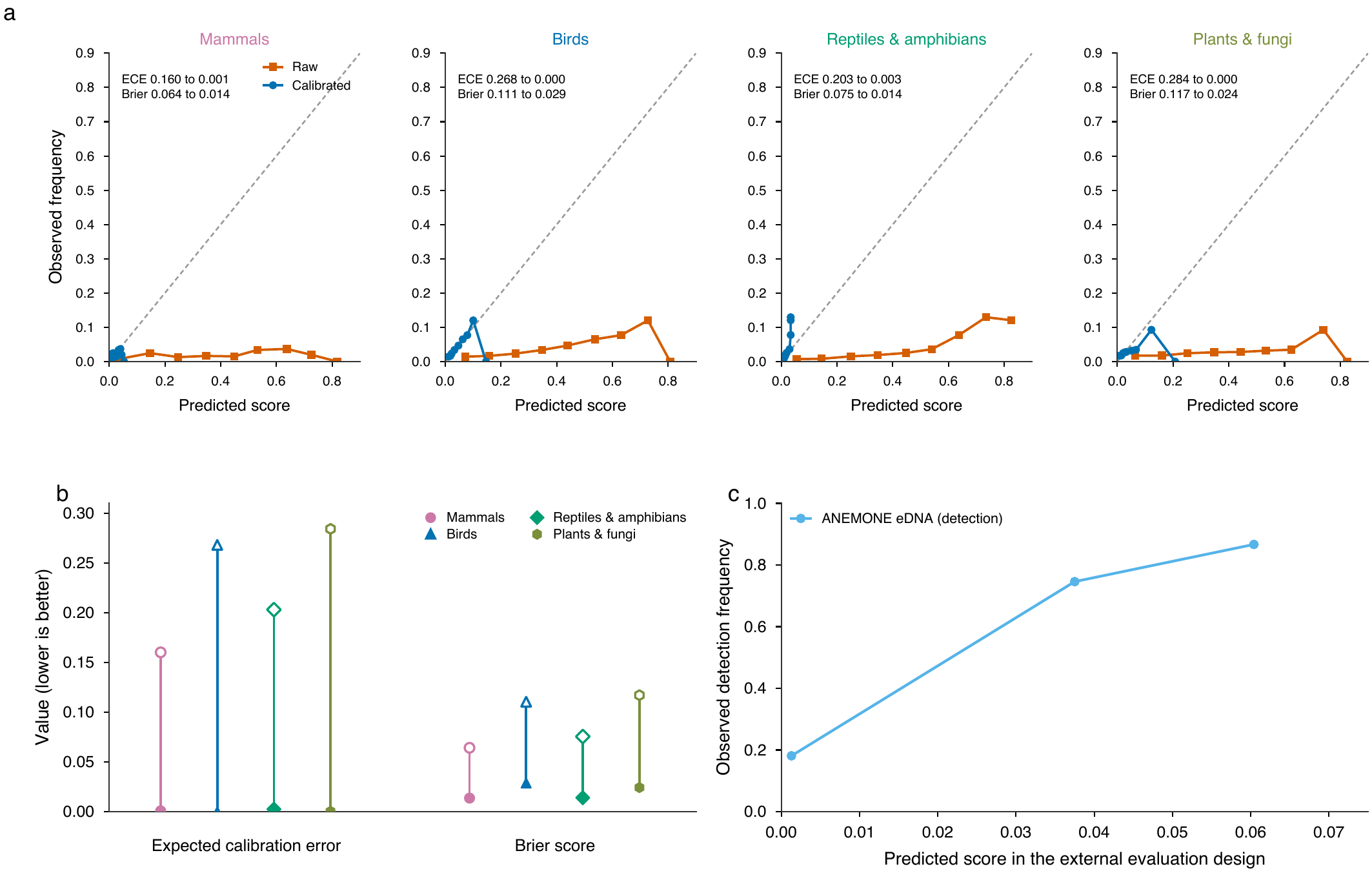

**Supplementary Fig. 5 | Calibration of model scores under internal and external observation designs.** a, Reliability curves before and after isotonic calibration in the spatially separated internal experiment for mammals, birds, reptiles and amphibians, and plants and fungi. b, Expected calibration error and Brier score before and after calibration; lower values indicate better calibration. c, External calibration against ANEMONE environmental-DNA detection and non-detection data. Calibrated values correspond to observed frequencies under the relevant evaluation design and are not absolute occurrence or occupancy probabilities.

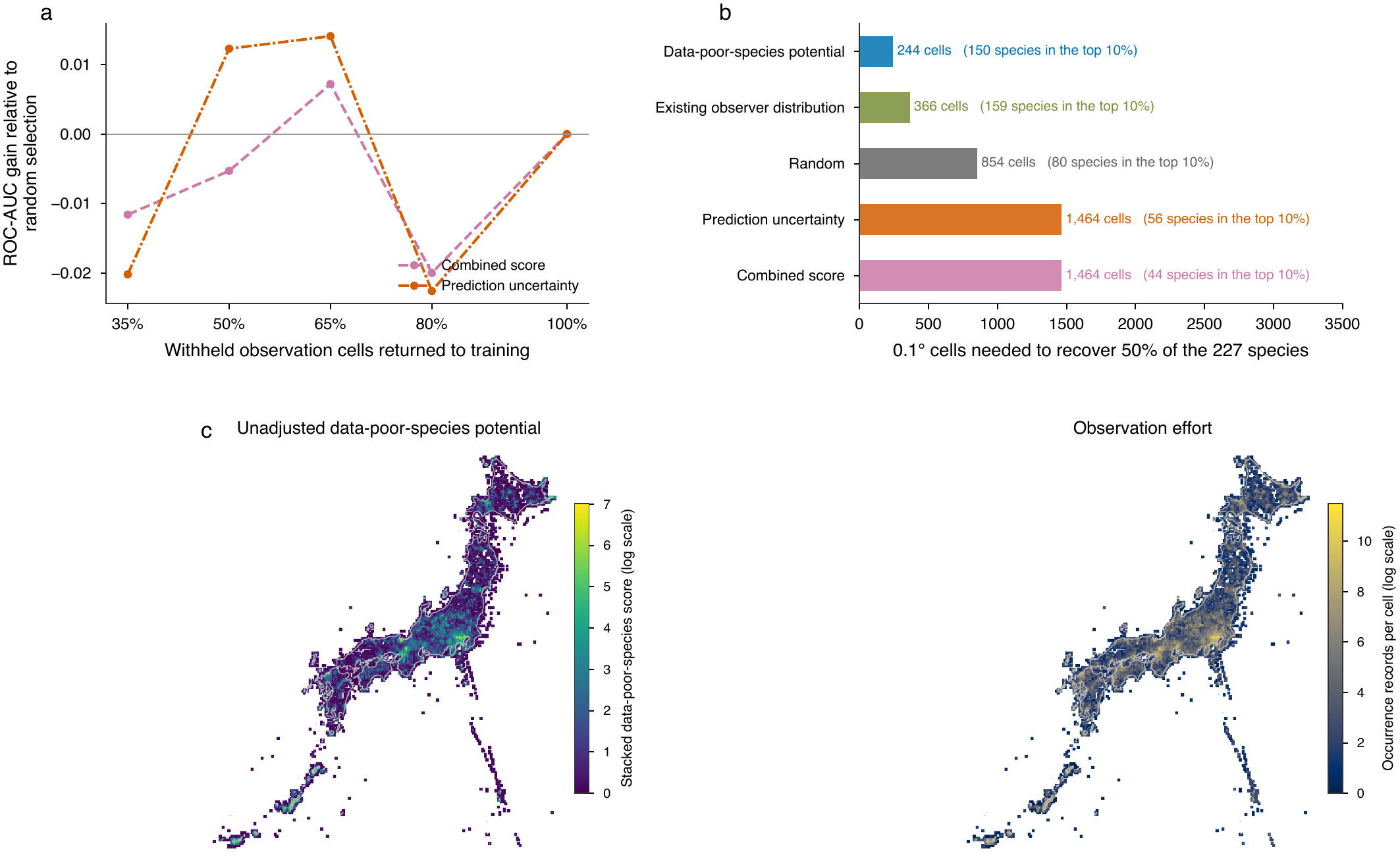

**Supplementary Fig. 6 | Evaluation of survey-priority strategies.** a, Gain in species-level ROC-AUC relative to random selection as withheld observation cells were returned to training. Values are means across two random seeds. b, Number of 0.1° cells required to recover 50% of 227 externally detected data-poor species; parentheses give the number detected at the saved 610-cell readout corresponding to the top 10% (the first evaluated budget point at or above the exact decile of 491 cells). c, National distributions of the unadjusted data-poor-species potential and observation effort. The independent discovery evaluation used the unadjusted 0.1° potential, not the residualized approximately 500-m layer deployed in the web application.

**Supplementary Table 1. Observation data sources, taxonomic composition, species-selection criteria and deployment routes.**

**(a) Observation data sources**

| **Data source** | **Observation period** | **Records in the canonical store** | **Records within the analysis window** | **Unique taxa within the analysis window** | **Data type** | **Notes** |
| --- | --- | --- | --- | --- | --- | --- |
| Global Biodiversity Information Facility (GBIF) | 1990 onwards (analysis window 2020–2025) | 668,280 | 290,686 | 3,878 | Presence-only | Records carrying this source tag; a record may carry more than one tag, so tagged counts do not sum to the integrated total. |
| iNaturalist | 1990 onwards (analysis window 2020–2025) | 1,176,211 | 929,729 | 26,710 | Presence-only | Records carrying this source tag; a record may carry more than one tag, so tagged counts do not sum to the integrated total. |
| eBird | 1990 onwards (analysis window 2020–2025) | 558,862 | 291,784 | 565 | Presence-only | Records carrying this source tag; a record may carry more than one tag, so tagged counts do not sum to the integrated total. |
| Ministry of the Environment Ikimono Log | 1990 onwards (analysis window 2020–2025) | Not separately tagged | Not separately tagged | Not separately tagged | Presence-only | Not resolvable as a distinct source tag in the canonical store; records reaching the store through aggregator publication are counted under the aggregator. |
| Researcher field surveys | 1990 onwards (analysis window 2020–2025) | Not separately tagged | Not separately tagged | Not separately tagged | Presence-only and structured survey | As above. |
| Integrated dataset | 1990 onwards (analysis window 2020–2025) | 2,322,578 | 1,461,834 | 26,869 | Integrated, deduplicated | 37,995 species over the whole store. Records with the same species, date and location were merged across sources, so source-level counts exceed the integrated total. |

**(b) Taxonomic composition and deployment**

| **Taxonomic group** | **Core species (≥20 records)** | **Species added beyond the core roster** | **Final deployed species** | **Deployed on the shared deep model** | **Deployed on a per-species ensemble** | **Deployed on the rare-species fallback** | **Long-tail pooling** | **Nominal minimum records** | **Effective minimum records in the deployed roster** | **External calibration anchor** |
| --- | --- | --- | --- | --- | --- | --- | --- | --- | --- | --- |
| Mammals | 52 | 9 | 61 | 0 | 61 | 0 | Not applied | 20 | 15 | None |
| Birds | 453 | 18 | 471 | 451 | 0 | 20 | Not applied | 20 | 15 | Satoyama bird route census (occupancy-based) |
| Reptiles & Amphibians | 98 | 14 | 112 | 0 | 112 | 0 | Not applied | 20 | 15 | None |
| Fishes | 229 | 251 | 480 | 480 | 0 | 0 | Applied | 5 | 1 | ANEMONE environmental DNA |
| Arthropods | 2,376 | 1,249 | 3,625 | 3,578 | 0 | 47 | Applied | 5 | 8 | None |
| Other invertebrates | 151 | 50 | 201 | 151 | 0 | 50 | Not applied | 20 | 15 | None |
| Plants & Fungi | 2,249 | 1,098 | 3,347 | 3,327 | 0 | 20 | Applied | 5 | 8 | Complete forest inventory |
| Total | 5,608 | 2,689 | 8,297 | 7,987 | 173 | 137 | Applied to three groups | NA | NA | NA |

Notes: the canonical store covers records from 1990 onwards; the analysis window applied to model training and evaluation is 2020–2025. Source-tagged counts do not sum to the integrated total because a record may carry more than one tag and duplicates were merged across sources. Core species are those with at least 20 records in the analysis window.

Abbreviations: AUC, area under the receiver operating characteristic curve; CI, confidence interval; ECE, expected calibration error; GNN, graph neural network; IQR, interquartile range; MLP, multilayer perceptron; NA, not applicable; TOST, two one-sided tests.

**Supplementary Table 2. Environmental covariates used in the final deployed models (41 nominal input columns).**

| **variable_id** | **variable_name** | **category** | **source_dataset** | **unit** | **native_resolution** | **temporal_coverage** | **effective** | **note** |
| --- | --- | --- | --- | --- | --- | --- | --- | --- |
| V01 | Annual total precipitation | Climate | National Land Numerical Information, Climatic Normals Mesh (G02; 1-km mesh) | mm | 1 km mesh (third-order mesh) | 1981-2010 climatological normals | Yes |  |
| V02 | Annual mean air temperature | Climate | National Land Numerical Information, Climatic Normals Mesh (G02; 1-km mesh) | degrees C | 1 km mesh (third-order mesh) | 1981-2010 climatological normals | Yes |  |
| V03 | Minimum air temperature in January | Climate | National Land Numerical Information, Climatic Normals Mesh (G02; 1-km mesh) | degrees C | 1 km mesh (third-order mesh) | 1981-2010 climatological normals | Yes |  |
| V04 | Maximum air temperature in August | Climate | National Land Numerical Information, Climatic Normals Mesh (G02; 1-km mesh) | degrees C | 1 km mesh (third-order mesh) | 1981-2010 climatological normals | Yes |  |
| V05 | Annual air temperature range (annual maximum minus annual minimum of monthly extremes) | Climate | National Land Numerical Information, Climatic Normals Mesh (G02; 1-km mesh) | degrees C | 1 km mesh (third-order mesh) | 1981-2010 climatological normals | Yes |  |
| V06 | Maximum snow depth (annual maximum of monthly deepest snow) | Climate | National Land Numerical Information, Climatic Normals Mesh (G02; 1-km mesh) | cm | 1 km mesh (third-order mesh) | 1981-2010 climatological normals | Yes |  |
| V07 | Annual sunshine duration | Climate | National Land Numerical Information, Climatic Normals Mesh (G02; 1-km mesh) | h | 1 km mesh (third-order mesh) | 1981-2010 climatological normals | Yes |  |
| V08 | Mean elevation within the 1-km mesh | Elevation | National Land Numerical Information, Elevation and Slope Angle Tertiary Mesh (G04-a-11) | m | 1 km mesh (third-order mesh) | 2011 edition (content reference date 2009-05-01) | Yes |  |
| V09 | Maximum elevation within the 1-km mesh | Elevation | National Land Numerical Information, Elevation and Slope Angle Tertiary Mesh (G04-a-11) | m | 1 km mesh (third-order mesh) | 2011 edition (content reference date 2009-05-01) | Yes |  |
| V10 | Minimum elevation within the 1-km mesh | Elevation | National Land Numerical Information, Elevation and Slope Angle Tertiary Mesh (G04-a-11) | m | 1 km mesh (third-order mesh) | 2011 edition (content reference date 2009-05-01) | Yes |  |
| V11 | Mean slope angle within the 1-km mesh | Slope and terrain | National Land Numerical Information, Elevation and Slope Angle Tertiary Mesh (G04-a-11) | degrees | 1 km mesh (third-order mesh) | 2011 edition (content reference date 2009-05-01) | Yes |  |
| V12 | Maximum slope angle within the 1-km mesh | Slope and terrain | National Land Numerical Information, Elevation and Slope Angle Tertiary Mesh (G04-a-11) | degrees | 1 km mesh (third-order mesh) | 2011 edition (content reference date 2009-05-01) | Yes |  |
| V13 | Forest stand volume per hectare (measured) | Forest resources | Forest resource aggregation mesh, 20 m (three prefectures under the Forest GIS Forum v2.0 specification; three prefectures from prefecture-specific open data, formal dataset names NA) | m3/ha | 20 m mesh | 2008-2022 (measurement year missing for 19.4% of cells) | No | Excluded from the effective set: the grid join used exact coordinate matching and did not snap to the 20 m grid, leaving a non-missing rate of 0.03-0.34% at training nodes; missing values were imputed with a constant (0) and permutation importance was at noise level. |
| V14 | Total forest stand volume per cell | Forest resources | Forest resource aggregation mesh, 20 m (three prefectures under the Forest GIS Forum v2.0 specification; three prefectures from prefecture-specific open data, formal dataset names NA) | m3 | 20 m mesh | 2008-2022 (measurement year missing for 19.4% of cells) | No | Excluded from the effective set: the grid join used exact coordinate matching and did not snap to the 20 m grid, leaving a non-missing rate of 0.03-0.34% at training nodes; missing values were imputed with a constant (0) and permutation importance was at noise level. |
| V15 | Mean tree height | Forest resources | Forest resource aggregation mesh, 20 m (three prefectures under the Forest GIS Forum v2.0 specification; three prefectures from prefecture-specific open data, formal dataset names NA) | m | 20 m mesh | 2008-2022 (measurement year missing for 19.4% of cells) | No | Excluded from the effective set: the grid join used exact coordinate matching and did not snap to the 20 m grid, leaving a non-missing rate of 0.03-0.34% at training nodes; missing values were imputed with a constant (0) and permutation importance was at noise level. |
| V16 | Stem density | Forest resources | Forest resource aggregation mesh, 20 m (three prefectures under the Forest GIS Forum v2.0 specification; three prefectures from prefecture-specific open data, formal dataset names NA) | stems/ha | 20 m mesh | 2008-2022 (measurement year missing for 19.4% of cells) | No | Excluded from the effective set: the grid join used exact coordinate matching and did not snap to the 20 m grid, leaving a non-missing rate of 0.03-0.34% at training nodes; missing values were imputed with a constant (0) and permutation importance was at noise level. |
| V17 | Slope angle (forest-mesh attribute) | Forest resources | Terrain attribute distributed with the 20 m forest resource mesh (not a forest measurement) | degrees | 20 m mesh | 2008-2022 (measurement year missing for 19.4% of cells) | No | Excluded from the effective set: the grid join used exact coordinate matching and did not snap to the 20 m grid, leaving a non-missing rate of 0.03-0.34% at training nodes; missing values were imputed with a constant (0) and permutation importance was at noise level. Terrain attribute carried by the forest mesh, not a forest resource measurement; elevation and slope are also supplied at 1 km by G04-a (V08-V12). |
| V18 | Elevation (forest-mesh attribute) | Forest resources | Terrain attribute distributed with the 20 m forest resource mesh (not a forest measurement) | m | 20 m mesh | 2008-2022 (measurement year missing for 19.4% of cells) | No | Excluded from the effective set: the grid join used exact coordinate matching and did not snap to the 20 m grid, leaving a non-missing rate of 0.03-0.34% at training nodes; missing values were imputed with a constant (0) and permutation importance was at noise level. Terrain attribute carried by the forest mesh, not a forest resource measurement; elevation and slope are also supplied at 1 km by G04-a (V08-V12). |
| V19 | Predicted mean diameter at breast height | Forest resources | Forest resource aggregation mesh, 20 m (three prefectures under the Forest GIS Forum v2.0 specification; three prefectures from prefecture-specific open data, formal dataset names NA) | cm | 20 m mesh | 2008-2022 (measurement year missing for 19.4% of cells) | No | Excluded from the effective set: the grid join used exact coordinate matching and did not snap to the 20 m grid, leaving a non-missing rate of 0.03-0.34% at training nodes; missing values were imputed with a constant (0) and permutation importance was at noise level. Model output rather than a field measurement; the column name of the source field is 'predicted'. |
| V20 | Estimated stand age | Forest resources | Forest resource aggregation mesh, 20 m (three prefectures under the Forest GIS Forum v2.0 specification; three prefectures from prefecture-specific open data, formal dataset names NA) | years | 20 m mesh | 2008-2022 (measurement year missing for 19.4% of cells) | No | Excluded from the effective set: the grid join used exact coordinate matching and did not snap to the 20 m grid, leaving a non-missing rate of 0.03-0.34% at training nodes; missing values were imputed with a constant (0) and permutation importance was at noise level. Model output rather than a field measurement; the column name of the source field is 'predicted'. |
| V21 | Mean annual increment | Forest resources | Forest resource aggregation mesh, 20 m (three prefectures under the Forest GIS Forum v2.0 specification; three prefectures from prefecture-specific open data, formal dataset names NA) | m3/ha/yr | 20 m mesh | 2008-2022 (measurement year missing for 19.4% of cells) | No | Excluded from the effective set: the grid join used exact coordinate matching and did not snap to the 20 m grid, leaving a non-missing rate of 0.03-0.34% at training nodes; missing values were imputed with a constant (0) and permutation importance was at noise level. Derived from the modelled stand age / a static unit price rather than measured directly. |
| V22 | Predicted stand volume per hectare (growth model) | Forest resources | Forest resource aggregation mesh, 20 m (three prefectures under the Forest GIS Forum v2.0 specification; three prefectures from prefecture-specific open data, formal dataset names NA) | m3/ha | 20 m mesh | 2008-2022 (measurement year missing for 19.4% of cells) | No | Excluded from the effective set: the grid join used exact coordinate matching and did not snap to the 20 m grid, leaving a non-missing rate of 0.03-0.34% at training nodes; missing values were imputed with a constant (0) and permutation importance was at noise level. Despite the source column name, 99.998% of values are identical to the measured stand volume. |
| V23 | Estimated forest resource value per hectare | Forest resources | Forest resource aggregation mesh, 20 m (three prefectures under the Forest GIS Forum v2.0 specification; three prefectures from prefecture-specific open data, formal dataset names NA) | JPY/ha | 20 m mesh | 2008-2022 (measurement year missing for 19.4% of cells) | No | Excluded from the effective set: the grid join used exact coordinate matching and did not snap to the 20 m grid, leaving a non-missing rate of 0.03-0.34% at training nodes; missing values were imputed with a constant (0) and permutation importance was at noise level. Derived from the modelled stand age / a static unit price rather than measured directly. |
| V24 | Proportion of natural land cover (forest and other natural land) within a 500 m radius | Land-use composition | Composite 20-m land-use layer built from national open datasets: agricultural parcel ('fude') polygons (2022), urban zoning areas (National Land Numerical Information A29), the national vegetation survey map 2024 (Biodiversity Center, Ministry of the Environment) and a 20-m forest resource mesh | % | 20 m (composite grid) | 2022 (parcel polygons) / 2024 (vegetation map); zoning A29-19 (2019 edition) | Yes |  |
| V25 | Proportion of agricultural land (paddy and cropland) within a 500 m radius | Land-use composition | Composite 20-m land-use layer built from national open datasets: agricultural parcel ('fude') polygons (2022), urban zoning areas (National Land Numerical Information A29), the national vegetation survey map 2024 (Biodiversity Center, Ministry of the Environment) and a 20-m forest resource mesh | % | 20 m (composite grid) | 2022 (parcel polygons) / 2024 (vegetation map); zoning A29-19 (2019 edition) | Yes |  |
| V26 | Proportion of built-up land within a 500 m radius | Land-use composition | Composite 20-m land-use layer built from national open datasets: agricultural parcel ('fude') polygons (2022), urban zoning areas (National Land Numerical Information A29), the national vegetation survey map 2024 (Biodiversity Center, Ministry of the Environment) and a 20-m forest resource mesh | % | 20 m (composite grid) | 2022 (parcel polygons) / 2024 (vegetation map); zoning A29-19 (2019 edition) | Yes |  |
| V27 | Proportion of open water within a 500 m radius | Land-use composition | Composite 20-m land-use layer built from national open datasets: agricultural parcel ('fude') polygons (2022), urban zoning areas (National Land Numerical Information A29), the national vegetation survey map 2024 (Biodiversity Center, Ministry of the Environment) and a 20-m forest resource mesh | % | 20 m (composite grid) | 2022 (parcel polygons) / 2024 (vegetation map); zoning A29-19 (2019 edition) | Yes |  |
| V28 | Mean vegetation naturalness within a 500 m radius | Land-use composition | National vegetation survey map 2024 (Biodiversity Center, Ministry of the Environment), vegetation-naturalness attribute, via the 20-m land-use composite | dimensionless | 20 m (composite grid) | 2024 (vegetation survey map 2024) | No | Excluded from the effective set: the mean was computed without excluding sentinel codes (open water / built-up areas), so the value is not interpretable as a naturalness index. |
| V29 | Distance to the nearest open-water cell | Distance to land-use classes | Derived from the 20-m land-use composite (see V24-V27) | m | 20 m (composite grid) | 2022 (parcel polygons) / 2024 (vegetation map); zoning A29-19 (2019 edition) | Yes |  |
| V30 | Distance to the nearest agricultural-land cell | Distance to land-use classes | Derived from the 20-m land-use composite (see V24-V27) | m | 20 m (composite grid) | 2022 (parcel polygons) / 2024 (vegetation map); zoning A29-19 (2019 edition) | Yes |  |
| V31 | Distance to the nearest built-up cell | Distance to land-use classes | Derived from the 20-m land-use composite (see V24-V27) | m | 20 m (composite grid) | 2022 (parcel polygons) / 2024 (vegetation map); zoning A29-19 (2019 edition) | Yes |  |
| V32 | Distance to the nearest coastline | Distance to other features | Natural Earth, Admin 0 - Countries (1:10m, v5 series, public domain); single polygon for Japan | m | 1:10m generalized coastline (6,952 vertices nationwide) | v5 series (patch release NA (not recoverable)) | Yes |  |
| V33 | Distance to the nearest river | Distance to other features | National Land Numerical Information, Rivers (W05; prefectural editions) | m | Vector (polyline/polygon vertices) | 2006-2009 (prefectural editions) | Yes (partial coverage) | Partial coverage: prefecture-level source data could not be retrieved for 29 of 47 prefectures; non-missing rate 0.4671 at the training nodes of the deployed core bird model. |
| V34 | Distance to the nearest lake or pond | Distance to other features | National Land Numerical Information, Lakes and Ponds (W09) | m | Vector (polyline/polygon vertices) | 2005 edition | Yes |  |
| V35 | Distance to the nearest railway line | Distance to other features | National Land Numerical Information, Railways (N02) | m | Vector (polyline/polygon vertices) | 2023 edition | Yes |  |
| V36 | Distance to the nearest city park | Distance to other features | National Land Numerical Information, City Parks (P13) | m | Vector (polyline/polygon vertices) | 2011 edition | Yes |  |
| V37 | Soil clay content (0-30 cm depth average) | Soil | SoilGrids 2.0 (ISRIC - World Soil Information), global soil property grids | % | 250 m global raster, queried at 0.1-degree grid points | 2020 (SoilGrids 2.0 release) | Yes |  |
| V38 | Soil sand content (0-30 cm depth average) | Soil | SoilGrids 2.0 (ISRIC - World Soil Information), global soil property grids | % | 250 m global raster, queried at 0.1-degree grid points | 2020 (SoilGrids 2.0 release) | Yes |  |
| V39 | Soil silt content (0-30 cm depth average) | Soil | SoilGrids 2.0 (ISRIC - World Soil Information), global soil property grids | % | 250 m global raster, queried at 0.1-degree grid points | 2020 (SoilGrids 2.0 release) | Yes |  |
| V40 | Hydrologic soil group (USDA classes A-D approximated from clay and sand content, ordinal-encoded) | Soil | SoilGrids 2.0 (ISRIC - World Soil Information), global soil property grids | dimensionless (ordinal) | 250 m global raster, queried at 0.1-degree grid points | 2020 (SoilGrids 2.0 release) | Yes |  |
| V41 | Summer NDVI (June-September median composite) | Satellite vegetation index | MODIS MOD13Q1 v061 16-day NDVI (NASA LP DAAC) | dimensionless | 250 m, sampled at 0.05-degree grid points | June-September 2023 (median composite) | Yes |  |

Notes: all 41 nominal input columns are listed individually with their data sources, units, native resolutions and temporal coverage. Preprocessing, missing-value imputation and standardization procedures are summarized in Supplementary Methods 1.4; implementation details of the feature pipeline and data-provider internal identifiers are not disclosed. Eleven forest-resource columns and one land-use column entered the input matrix but did not contribute to the reported results (see the ‘effective’ and ‘note’ columns); all reported performance was therefore obtained without forest-structure information. The distance-to-river column has partial spatial coverage (missing in 29 of 47 prefectures). Cells marked ‘NA (not recoverable)’ denote provenance that cannot be reconstructed: the input-version record was implemented after the deployed models were fitted.

Abbreviations: AUC, area under the receiver operating characteristic curve; CI, confidence interval; ECE, expected calibration error; GNN, graph neural network; IQR, interquartile range; MLP, multilayer perceptron; NA, not applicable; TOST, two one-sided tests.

**Supplementary Table 3. Deployment route and species-selection settings by taxonomic group.**

| **Taxonomic group** | **Deployed model family** | **Rare-species fallback in the deployed set** | **Core threshold (records)** | **Long-tail threshold (records)** | **Deployed species** | **Score calibration applied before deployment** |
| --- | --- | --- | --- | --- | --- | --- |
| Mammals | Per-species ensemble | No | 20 | Not applicable | 61 | Yes |
| Birds | Shared-representation deep model | Yes | 20 | Not applicable | 471 | Yes |
| Reptiles & Amphibians | Per-species ensemble | No | 20 | Not applicable | 112 | Yes |
| Fishes | Shared-representation deep model | No | 20 | 5 | 480 | Yes |
| Arthropods | Shared-representation deep model | Yes | 20 | 5 | 3,625 | Yes |
| Other invertebrates | Shared-representation deep model | Yes | 20 | Not applicable | 201 | Yes |
| Plants & Fungi | Shared-representation deep model | Yes | 20 | 5 | 3,347 | Yes |

Notes: Model architectures, training settings and hyperparameters are described in SM1.6, SM1.8 and SM1.9. The rare-species fallback denotes deployed species predicted by a per-species ensemble outside the taxon core roster.

Abbreviations: AUC, area under the receiver operating characteristic curve; CI, confidence interval; ECE, expected calibration error; GNN, graph neural network; IQR, interquartile range; MLP, multilayer perceptron; NA, not applicable; TOST, two one-sided tests.

**Supplementary Table 4. Statistical summaries of model comparisons, long-tail pooling and ablation analyses.**

**(a) Shared deep model versus per-species random forest**

| **Taxonomic group** | **Evaluable species** | **Median AUC, shared model** | **Median AUC, species-specific random forest** | **Median paired difference** | **95% CI** | **Raw P** | **Holm-adjusted P** | **Test** | **Species with higher shared-model AUC** |
| --- | --- | --- | --- | --- | --- | --- | --- | --- | --- |
| Mammals | 11 | 0.619 | 0.629 | -0.027 | [-0.181, 0.069] | 0.320 | 0.577 | Wilcoxon signed-rank on paired species AUC | 4/11 |
| Birds | 120 | 0.757 | 0.707 | 0.025 | [0.012, 0.036] | <0.001 | 0.003 | Wilcoxon signed-rank on paired species AUC | 77/120 |
| Reptiles & amphibians | 30 | 0.647 | 0.69 | -0.021 | [-0.045, -0.004] | 0.064 | 0.191 | Wilcoxon signed-rank on paired species AUC | 9/30 |
| Fishes | 98 | 0.598 | 0.539 | 0.04 | [0.001, 0.086] | 0.015 | 0.076 | Wilcoxon signed-rank on paired species AUC | 59/98 |
| Arthropods | 97 | 0.593 | 0.5 | 0.078 | [0.057, 0.118] | <0.001 | <0.001 | Wilcoxon signed-rank on paired species AUC | 68/97 |
| Other invertebrates | 32 | 0.621 | 0.535 | 0.148 | [0.003, 0.230] | 0.022 | 0.086 | Wilcoxon signed-rank on paired species AUC | 21/32 |
| Plants & fungi | 114 | 0.559 | 0.514 | 0.019 | [-0.025, 0.045] | 0.288 | 0.577 | Wilcoxon signed-rank on paired species AUC | 59/114 |

**(b) Benchmark comparisons**

| **Analysis set** | **Model** | **Evaluable species** | **Median AUC** | **Reference model** | **Paired difference** | **P** | **Notes** |
| --- | --- | --- | --- | --- | --- | --- | --- |
| Deployment pipeline, complete forest inventory | Shared representation | 125 | 0.832 | NA | NA | NA | Predictor sets differed among models. 59 plots, 125 species; distinct from the 122-species, 60-plot external validation in ST5b. |
| Deployment pipeline, complete forest inventory | MaxEnt | 125 | 0.806 | Shared representation | -0.03 | 0.005 | Predictor sets differed among models. |
| Deployment pipeline, complete forest inventory | Random forest | 125 | 0.761 | Shared representation | -0.045 | <0.001 | Predictor sets differed among models. |
| Fully controlled comparison | MaxEnt | 28 | 0.556 | NA | NA | NA | Identical occurrence records, background locations, environmental covariates and spatial splits. |
| Fully controlled comparison | Random forest | 28 | 0.55 | NA | NA | NA | Identical occurrence records, background locations, environmental covariates and spatial splits. |
| Fully controlled comparison | MLP | 28 | 0.532 | NA | NA | NA | Identical occurrence records, background locations, environmental covariates and spatial splits. |
| Fully controlled comparison | GNN | 28 | 0.536 | NA | NA | NA | Identical occurrence records, background locations, environmental covariates and spatial splits. |

**(c) Long-tail pooling**

| **Taxonomic group** | **External evaluation** | **Evaluable species before** | **Evaluable species after** | **Median AUC before** | **Median AUC after** | **Evaluation frame before** | **Evaluation frame after** | **Species evaluable in both arms** | **Median paired difference** | **95% CI** | **P** | **Equivalence result (±0.01 AUC, TOST, internal holdout)** |
| --- | --- | --- | --- | --- | --- | --- | --- | --- | --- | --- | --- | --- |
| Fishes | ANEMONE MiFish environmental DNA | 144 | 200 | 0.683 | 0.724 | 148 sites | 150 sites | 144 | 0.0122 | [+0.0062, +0.0228] | <0.001 | Not equivalent |
| Arthropods | Butterfly route-census transects | 45 | 45 | 0.684 | 0.733 | 33 sites | 32 sites | 45 | 0.0376 | [+0.0157, +0.0704] | 0.001 | Equivalent |
| Plants & fungi | Complete forest inventory | 122 | 125 | 0.837 | 0.832 | 59 plots (common frame) | 59 plots (common frame) | 122 | 0.0009 | [-0.0092, +0.0085] | 0.524 | Not tested |
| Fishes | ANEMONE MiFish environmental DNA | 120 | 120 | 0.703 | 0.726 | site denominator unchanged | site denominator unchanged | 120 | 0.0064 | [-0.0023, +0.0125] | 0.169 | Not applicable |

**(c continued) Species that became evaluable only after pooling**

| **Taxonomic group** | **External evaluation** | **Newly evaluable species** | **Median AUC** | **Interquartile range** | **Fraction below chance** | **Median AUC of the previously evaluable species in the same run** |
| --- | --- | --- | --- | --- | --- | --- |
| Fishes | ANEMONE MiFish environmental DNA | 56 | 0.715 | 0.645-0.815 | 0.018 | 0.726 |

**(c-iii) Effect of the score column on the long-tail after-arms**

| **Taxonomic group** | **External evaluation** | **Evaluable species** | **Scored rows** | **Median AUC, uncalibrated score** | **Median AUC, calibrated score** | **Difference attributable to the score column** | **Distinct scores, uncalibrated** | **Distinct scores, calibrated** | **Largest tied group, uncalibrated** | **Largest tied group, calibrated** | **Artefact, uncalibrated** | **Artefact, calibrated** | **Notes** |
| --- | --- | --- | --- | --- | --- | --- | --- | --- | --- | --- | --- | --- | --- |
| Plants & fungi | Complete forest inventory | 125 | 7,375 | 0.8511 | 0.8322 | -0.0189 | 1,316 | 21 | 211 | 2,358 | eval_biodic_plantmin5 | eval_biodic_final | Same species and same scored rows in both columns; only the score differs. Both before-arms use the calibrated column: the evaluation code used for the before-arms reads prob, and prob was already the calibrated column by then; for plants and fungi this was confirmed directly on 2026-08-08 by reconstructing the before-arm cell values from the retained prediction surface (r = 0.9985, MAE 0.0043 against the published per-species file). The reported plants contrast in panel (c) is paired on the common 59-plot frame with the calibrated column in both arms; the like-for-like calibrated fish comparison is 0.683 to 0.724. |
| Fishes | ANEMONE MiFish environmental DNA | 200 | 30,000 | 0.7355 | 0.7237 | -0.0118 | 2,632 | 54 | 1,290 | 15,745 | eval_edna_min5 | eval_edna_deployed | Same species and same scored rows in both columns; only the score differs. Both before-arms use the calibrated column: the evaluation code used for the before-arms reads prob, and prob was already the calibrated column by then; for plants and fungi this was confirmed directly on 2026-08-08 by reconstructing the before-arm cell values from the retained prediction surface (r = 0.9985, MAE 0.0043 against the published per-species file). The reported plants contrast in panel (c) is paired on the common 59-plot frame with the calibrated column in both arms; the like-for-like calibrated fish comparison is 0.683 to 0.724. |

**(d) Ablation analyses**

| **Comparison** | **Architecture** | **Taxonomic group** | **Evaluable species** | **Effect estimate (ROC-AUC)** | **95% CI** | **P** | **Notes** |
| --- | --- | --- | --- | --- | --- | --- | --- |
| Shared versus individual representation | GNN | Mammals | 25 | 0.032 | NA | NA | Positive values favour the shared model. P from the species-clustered Wilcoxon test (SM2.5); runs without per-record data carry no P. |
| Shared versus individual representation | GNN | Mammals | 24 | 0.0053 | [-0.0116, 0.0195] | 0.107 | Positive values favour the shared model. P from the species-clustered Wilcoxon test (SM2.5); runs without per-record data carry no P. |
| Shared versus individual representation | GNN | Mammals | 24 | -0.0006 | [-0.0176, 0.0143] | 0.491 | Positive values favour the shared model. P from the species-clustered Wilcoxon test (SM2.5); runs without per-record data carry no P. |
| Shared versus individual representation | GNN | Birds | 25 | 0.011 | NA | NA | Positive values favour the shared model. P from the species-clustered Wilcoxon test (SM2.5); runs without per-record data carry no P. |
| Shared versus individual representation | GNN | Birds | 25 | 0.0114 | [0.0081, 0.0148] | <0.001 | Positive values favour the shared model. P from the species-clustered Wilcoxon test (SM2.5); runs without per-record data carry no P. |
| Shared versus individual representation | GNN | Reptiles & amphibians | 25 | 0.0144 | [0.0077, 0.0234] | <0.001 | Positive values favour the shared model. P from the species-clustered Wilcoxon test (SM2.5); runs without per-record data carry no P. |
| Shared versus individual representation | GNN | Reptiles & amphibians | 25 | 0.0093 | [0.0045, 0.0205] | <0.001 | Positive values favour the shared model. P from the species-clustered Wilcoxon test (SM2.5); runs without per-record data carry no P. |
| Shared versus individual representation | GNN | Reptiles & amphibians | 25 | 0.0101 | [0.0041, 0.0183] | <0.001 | Positive values favour the shared model. P from the species-clustered Wilcoxon test (SM2.5); runs without per-record data carry no P. |
| Shared versus individual representation | GNN | Arthropods | 25 | 0.009 | NA | NA | Positive values favour the shared model. P from the species-clustered Wilcoxon test (SM2.5); runs without per-record data carry no P. |
| Shared versus individual representation | MLP | Birds | 25 | 0.0159 | [0.0135, 0.0209] | <0.001 | Positive values favour the shared model. P from the species-clustered Wilcoxon test (SM2.5); runs without per-record data carry no P. |
| Shared versus individual representation | MLP | Reptiles & amphibians | 25 | 0.0307 | [0.0179, 0.0435] | <0.001 | Positive values favour the shared model. P from the species-clustered Wilcoxon test (SM2.5); runs without per-record data carry no P. |
| Graph definition | GNN | Birds | 25 | 0.0 | [-0.0003, 0.0004] | 0.653 | distance-weighted edges versus proximity; equivalent to zero within ±0.01 (TOST). |
| Graph definition | GNN | Birds | 25 | -0.0017 | [-0.0024, -0.0008] | 0.003 | river-corridor edges versus proximity; equivalent to zero within ±0.01 (TOST). |
| Graph definition | GNN | Reptiles & amphibians | 25 | 0.0011 | [0.0001, 0.0021] | 0.005 | distance-weighted edges versus proximity; equivalent to zero within ±0.01 (TOST). |
| Graph definition | GNN | Reptiles & amphibians | 25 | -0.0181 | [-0.0288, -0.0065] | 0.001 | catchment edges versus weighted; not shown equivalent to zero within ±0.01 (TOST). |
| Graph definition | GNN | Fishes | 25 | 0.0027 | [0.0017, 0.0045] | <0.001 | distance-weighted edges versus proximity; equivalent to zero within ±0.01 (TOST). |
| Graph definition | GNN | Fishes | 25 | -0.0149 | [-0.0303, -0.0051] | 0.182 | catchment edges versus weighted; not shown equivalent to zero within ±0.01 (TOST). |
| Graph definition | GNN | Mammals | 23 | -0.0051 | [-0.0202, 0.0097] | 0.482 | connectivity-repaired graph versus sparse graph; not shown equivalent to zero within ±0.01 (TOST). The manipulation also relocates background cells: the graph-free random-forest control moves by +0.0154, so this contrast does not isolate the graph. |
| Graph definition | GNN | Reptiles & amphibians | 25 | -0.0027 | [-0.007, 0.0001] | 0.191 | connectivity-repaired graph versus sparse graph; equivalent to zero within ±0.01 (TOST). The manipulation also relocates background cells: the graph-free random-forest control moves by -0.0053, so this contrast does not isolate the graph. |
| Number of jointly trained species | GNN | Birds | 6 | -0.027 | [-0.0677, -0.0106] | 0.219 | 6 species trained jointly; reference is the species-specific random forest. |
| Number of jointly trained species | GNN | Birds | 6 | -0.0307 | [-0.0597, -0.0091] | 0.156 | 12 species trained jointly; reference is the species-specific random forest. |
| Number of jointly trained species | GNN | Birds | 6 | -0.0216 | [-0.0535, -0.0103] | 0.031 | 25 species trained jointly; reference is the species-specific random forest. |
| Number of jointly trained species | GNN | Birds | 6 | -0.0148 | [-0.0295, +0.0039] | 0.219 | 50 species trained jointly; reference is the species-specific random forest. |
| Number of jointly trained species | GNN | Birds | 6 | -0.0057 | [-0.0307, +0.0068] | 0.438 | 100 species trained jointly; reference is the species-specific random forest. |
| Number of jointly trained species | MLP | Birds | 6 | -0.0291 | [-0.0796, -0.0039] | 0.156 | 6 species trained jointly; reference is the species-specific random forest. |
| Number of jointly trained species | MLP | Birds | 6 | -0.0186 | [-0.0463, -0.0047] | 0.063 | 12 species trained jointly; reference is the species-specific random forest. |
| Number of jointly trained species | MLP | Birds | 6 | -0.0176 | [-0.0614, +0.0026] | 0.063 | 25 species trained jointly; reference is the species-specific random forest. |
| Number of jointly trained species | MLP | Birds | 6 | -0.0182 | [-0.0282, -0.0072] | 0.094 | 50 species trained jointly; reference is the species-specific random forest. |
| Number of jointly trained species | MLP | Birds | 6 | 0.001 | [-0.0321, +0.0142] | 0.844 | 100 species trained jointly; reference is the species-specific random forest. |

**(e) Non-spatial random versus spatial block partitioning**

| **Taxonomic group** | **Species evaluable under both designs** | **Median ROC-AUC, spatial block split** | **Median ROC-AUC, random split** | **Median inflation** | **P** |
| --- | --- | --- | --- | --- | --- |
| Mammals | 11 | 0.605 | 0.867 | 0.343 | <0.001 |
| Birds | 202 | 0.706 | 0.838 | 0.101 | <0.001 |
| Reptiles & Amphibians | 30 | 0.626 | 0.845 | 0.222 | <0.001 |
| Fishes | 75 | 0.599 | 0.916 | 0.323 | <0.001 |
| Arthropods | 123 | 0.568 | 0.806 | 0.21 | <0.001 |
| Other invertebrates | 23 | 0.631 | 0.907 | 0.269 | <0.001 |
| Plants & Fungi | 90 | 0.548 | 0.811 | 0.278 | <0.001 |
| All groups | 554 | 0.634 | 0.858 | 0.185 | NA |

Notes: AUC denotes the area under the receiver operating characteristic curve. P values in panel (a) are two-sided Wilcoxon signed-rank tests over species, adjusted across the seven groups by the Holm method. Panel (b) separates the deployment-pipeline comparison, in which predictor sets differed among models, from the fully controlled comparison, in which they did not. In panel (c) the plants-and-fungi contrast is paired on a common 59-plot frame and the calibrated score column in both arms; the fish and arthropod contrasts are unpaired across their stated frames, with the paired statistic restricted to species evaluable in both arms. In panel (d) effect estimates are medians of paired per-species differences on a common spatial holdout, confidence intervals are bootstrapped with the species as the resampling unit, and P values are two-sided Wilcoxon signed-rank tests over species (replicates from seeds and blocks are collapsed within species before testing). For the 'Number of jointly trained species' rows the test is over six species, so the smallest attainable two-sided P is 0.031, and these ten P values are not adjusted for multiplicity and are reported for completeness rather than as a set of independent tests; in those rows the effect estimate and the confidence interval summarise all paired records whereas the P value is computed on the six species means, so a confidence interval that excludes zero can accompany P > 0.05. TOST, two one-sided tests.

Abbreviations: AUC, area under the receiver operating characteristic curve; CI, confidence interval; ECE, expected calibration error; GNN, graph neural network; IQR, interquartile range; MLP, multilayer perceptron; NA, not applicable; TOST, two one-sided tests.

**Supplementary Table 5. External validation datasets and predictive performance.**

**(a) External validation datasets**

| **Dataset** | **Taxonomic coverage** | **Survey design** | **Total sites** | **Evaluable sites** | **Leakage-controlled evaluable sites** | **Evaluable species** | **Interpretation of non-records** |
| --- | --- | --- | --- | --- | --- | --- | --- |
| Certified natural sites | All seven groups | Certified-site species inventories | 393 | 348 | 335 | 1,302 | Pseudo-absence; candidate species of the same group not listed at the site |
| ANEMONE MiFish environmental DNA | Fishes | Replicated water sampling, metabarcoding | 148 | 147 | Not applicable | 144 | Non-detection; not confirmed absence (detection probability below one) |
| ANEMONE MiFish, occupancy-corrected subset | Fishes | As above, with a multispecies occupancy model | 137 | 123 | Not applicable | 126 | Detection–non-detection labels; occupancy used for the prevalence denominator only |
| Monitoring Sites 1000 | Mammals, birds, other invertebrates, plants | Structured monitoring | 109 | 106 | Not applicable | 102 | Non-record; may include imperfect detection |
| Complete forest inventory | Trees | Complete stem enumeration in survey plots | 60 | 59 | Not applicable | 122 | Confirmed absence: every stem in the plot was counted |
| National Forest Inventory | Trees | Standardized national forest plots | 15,835 | 15,835 | Not applicable | 270 | Non-record within standardized forest plots; not a confirmed absence, unlike the complete forest inventories |

**(b) External validation performance**

| **Dataset** | **Evaluation level** | **Evaluation units** | **Median ROC-AUC** | **Leakage-controlled median ROC-AUC** | **Additional metric** | **Analysis note** |
| --- | --- | --- | --- | --- | --- | --- |
| Certified natural sites | Site-level community | 348 | 0.894 | 0.871 | Recorded species sit in the top 16.3% of the same-group candidate list | Leakage control removes evaluation pairs with a same-species training record within 2 km. |
| ANEMONE MiFish environmental DNA | Site-level community | 147 | 0.815 | Not applicable | IQR 0.724–0.882 over 147 evaluable sites | Naive detection labels. The 147-site generation; the file holds 148 sites and one admits no AUC. |
| Monitoring Sites 1000 | Site-level community | 106 | 0.724 | Not applicable | IQR 0.633–0.793 over 106 evaluable sites | Survey period does not overlap the model observation window. 109 sites in total; 106 admit an AUC. |
| Certified natural sites | Species-level | 1,302 | 0.672 | 0.650 (n = 1,224; IQR 0.488–0.817) | IQR 0.500–0.846 | Unadjusted evaluation. The leakage-controlled value is reported in the adjacent column and plotted in Supplementary Fig. 4b. |
| ANEMONE MiFish environmental DNA | Species-level | 144 | 0.683 | Not applicable | IQR 0.605–0.777 | Non-records have dataset-specific meanings. |
| ANEMONE MiFish, occupancy-corrected subset | Species-level | 126 | 0.776 | Not applicable | IQR 0.701–0.877 | Occupancy-corrected subset; different species and site population from the detection evaluation, so the two are not a paired contrast. |
| Monitoring Sites 1000 | Species-level | 102 | 0.69 | Not applicable | IQR 0.605–0.801 | Non-records have dataset-specific meanings. |
| National Forest Inventory | Species-level | 270 | 0.782 | Not applicable | IQR 0.693–0.857 | Non-records are non-records within standardized plots, not confirmed absences. |
| Complete forest inventory | Species-level | 122 | 0.841 | Not applicable | IQR 0.742–0.927 | Pre-tail-pooling core run, 60 plots and 122 species. Distinct from the 125-species, 59-plot deployment-pipeline comparison in ST4b. |
| Certified natural sites, species added by long-tail pooling | Species-level | 127 | 0.5 | Not applicable | IQR 0.429–0.887; 49.6% below 0.5 | Subset of the unadjusted certified-site evaluation above. Species present in the min_obs=5 shared roster but not in the core roster, for fishes, arthropods and plants and fungi. |
| Complete forest inventory | Site-level community | 59 | 0.755 | Not applicable | 60 plots | Pre-tail-pooling core; recovered from the archived pre-pooling evaluation. |

Notes: site-level community AUC evaluates whether species recorded at a site rank above candidate species of the same group that were not recorded there. Species-level AUC evaluates whether sites at which a species was recorded rank above sites at which it was not. The two are different evaluations and are not directly comparable. The meaning of a non-record differs among datasets, as given in panel (a).

Abbreviations: AUC, area under the receiver operating characteristic curve; CI, confidence interval; ECE, expected calibration error; GNN, graph neural network; IQR, interquartile range; MLP, multilayer perceptron; NA, not applicable; TOST, two one-sided tests.

**Supplementary Table 6. Calibration performance for internal spatial transfer and external validation datasets.**

**(a) Internal spatial calibration transfer**

| **Taxonomic group** | **Calibration region** | **Evaluation region** | **Test prevalence** | **ECE before** | **ECE after** | **Brier before** | **Brier after** | **Mean prediction before** | **Mean prediction after** |
| --- | --- | --- | --- | --- | --- | --- | --- | --- | --- |
| Mammals | Western third of the evaluation window | Eastern third, separated by a buffer | 0.0138 | 0.160 | 0.001 | 0.064 | 0.014 | 0.174 | 0.015 |
| Birds | Western third of the evaluation window | Eastern third, separated by a buffer | 0.0299 | 0.268 | 0.000 | 0.111 | 0.029 | 0.298 | 0.030 |
| Reptiles & amphibians | Western third of the evaluation window | Eastern third, separated by a buffer | 0.0143 | 0.203 | 0.003 | 0.075 | 0.014 | 0.217 | 0.012 |
| Plants & fungi | Western third of the evaluation window | Eastern third, separated by a buffer | 0.0248 | 0.284 | 0.000 | 0.117 | 0.024 | 0.309 | 0.025 |

**(b) External-data calibration**

| **Dataset** | **Taxonomic group** | **Calibration target** | **Spatial split** | **Evaluation cells** | **ECE before** | **ECE after** | **Brier before** | **Brier after** | **Notes** |
| --- | --- | --- | --- | --- | --- | --- | --- | --- | --- |
| Complete forest inventory | Trees | Presence–absence within survey plots | W->E | Not retained | 0.1753 | 0.03 | Not retained | Not retained | Quantile bins; not comparable with the equal-width bins used for the internal calibration. |
| Complete forest inventory | Trees | Presence–absence within survey plots | E->W | Not retained | 0.1807 | 0.0258 | Not retained | Not retained | Quantile bins; not comparable with the equal-width bins used for the internal calibration. |
| ANEMONE MiFish environmental DNA | Fishes | Detection–non-detection | W->E | 9,148 | 0.161 | 0.0343 | 0.1606 | 0.1112 | Detection, not occupancy: the calibration target is the probability of eDNA detection, and detection probability is below one. Reliability curve in Supplementary Fig. 5c. |
| ANEMONE MiFish environmental DNA | Fishes | Detection–non-detection | E->W | 8,941 | 0.217 | 0.0349 | 0.2165 | 0.1471 | Detection, not occupancy: the calibration target is the probability of eDNA detection, and detection probability is below one. Reliability curve in Supplementary Fig. 5c. |

Notes: ECE, expected calibration error. Internal calibration targets the frequency of presences among presence and background locations in the evaluation design and is not an absolute occurrence or occupancy probability. Bin definitions differ between the internal and external evaluations, so their ECE values are not directly comparable.

Abbreviations: AUC, area under the receiver operating characteristic curve; CI, confidence interval; ECE, expected calibration error; GNN, graph neural network; IQR, interquartile range; MLP, multilayer perceptron; NA, not applicable; TOST, two one-sided tests.

**Supplementary Table 7. Definitions, spatial units, intended uses and limitations of web-application outputs.**

| **Output** | **Definition** | **Spatial unit** | **Intended use** | **Main limitations** |
| --- | --- | --- | --- | --- |
| Existing occurrence records | Quality-controlled records located inside the region the user specified, shown separately from predictions. | Point records | Confirming what has already been recorded. | Locations of species of conservation concern are coarsened or withheld. |
| Candidate species list | Deployed species ranked by their calibrated prediction aggregated over the specified region. | Region aggregate | Selecting species to verify in the field. | A model candidate list, not a verified regional inventory. |
| Species-level prediction map | Calibrated relative occurrence score for one species. | 20 m output grid | Locating candidate habitat within a region. | 20 m is the output and display unit; it does not certify ecological accuracy at that resolution. |
| Biodiversity prediction map | Calibrated species scores stacked across the 8,290 deployed species remaining after seven human and domesticated taxa were removed, and adjusted for observation effort, then expressed relative to the selected region. Alien species are retained; native-only and alien-only layers are served alongside the all-species layer. | 20 m output grid, aggregated for display | Screening for areas of higher biodiversity potential. | Not an absolute species count or a sum of occupancy probabilities; values from different regions are not on a common scale. |
| Conservation-priority map | Calibrated species scores weighted by the inverse of each species' nationwide mean calibrated score, summed within each cell and expressed relative to the selected region. Native species only; including alien species leaves the ranking almost unchanged (Supplementary Methods 7.4). | 20 m output grid, aggregated for display | Screening for areas where species with lower nationwide mean calibrated scores overlap. | Not a comprehensive conservation priority: extinction risk, future land-conversion risk, cost, land tenure and manageability are not included. |
| Survey-priority map | Predicted concentration of data-poor species after accounting for existing observation effort, expressed relative to the specified region. | About 500 m local layer | Choosing where to survey next for data-poor species. | The external validation applies to the unadjusted 0.1° national potential; the deployed residual layer at about 500 m was not validated under the same design. |

Notes: calculation details for each output are reported at overview level only. All regional indices are expressed relative to the region the user specified, so values obtained for different regions are not on a common absolute scale. The 20 m grid and the approximately 500 m layer are output and display units and do not certify ecological accuracy at those resolutions.

Abbreviations: AUC, area under the receiver operating characteristic curve; CI, confidence interval; ECE, expected calibration error; GNN, graph neural network; IQR, interquartile range; MLP, multilayer perceptron; NA, not applicable; TOST, two one-sided tests.
